# A human neuronal model of sporadic Alzheimer’s disease induced by *FBXO2* downregulation shows Aβ aggregation, tau hyperphosphorylation and functional network impairment

**DOI:** 10.1101/2024.09.01.610673

**Authors:** Alicia González Díaz, Andrea Possenti, Gustavo Antonio Urrutia, Yuqi Bian, Shekhar Kedia, Dorothea Boeken, Christine M. Lim, Danilo Licastro, Benedetta Mannini, David Klenerman, Michele Vendruscolo

**Affiliations:** Centre for Misfolding Diseases, University of Cambridge, CB2 1EW Cambridge, UK; Yusuf Hamied Department of Chemistry, University of Cambridge, CB2 1EW Cambridge, UK; UK Dementia Research Institute, University of Cambridge, Cambridge, CB2 0AH, UK; AREA Science Park, Padriciano 99, 34149 Trieste, Italy

## Abstract

Sporadic Alzheimer’s disease (sAD) arises from a complex interplay between genetic and environmental factors that remains poorly understood, making it challenging to develop accurate cell models. To address this problem, by hypothesing that the early disease sAD states can be characterised by transcriptomic fingerprints, we assessed the effect on Aβ aggregation in human neuroblastoma cells a set of genes obtained by analysing snRNA-seq data from post-mortem AD patients. We then validated the most effective genes in human iPSC-derived cortical neurons, and selected *FBXO2*, a gene encoding a subunit of the ubiquitin protein ligase complex SCF, for further analysis. We found that early downregulation of *FBXO2* in human iPSC-derived cortical neurons resulted in Aβ aggregation, tau hyperphosphorylation, and structural and functional neuronal network impairment. Based on these results, we report a neuronal sAD model (*FBXO2* KD sAD) that recapitulates a set of molecular hallmarks of sAD. We suggest that this strategy can be expanded towards the generation of panels of preclinical stem cell-derived models that recapitulate the molecular complexity of the broad spectrum of AD patients.

## Introduction

Alzheimer’s disease (AD) is the most common cause of dementia, with projections indicating it will afflict over 100 million patients worldwide by 2050 (*1, 2*) unless effective treatments will become widely available (*3–6*). Familial Alzheimer’s disease (fAD) accounts for about 5% of cases, showing autosomal dominant inheritance and high degree of penetrance, as is generally caused by the presence of mutations in *APP*, which encodes for amyloid precursor protein (APP), and in *PSEN1* and *PSEN2*, which encode for presenilin-1 and presenilin-2, two enzymes responsible for the proteolytic cleavage of APP and the subsequent generation of amyloid-β peptides (Aβ) (*5, 7*). The remaining 95% of cases are sporadic (sAD), being the result of a complex interaction between genetic vulnerability and environmental triggers, including contact to toxic solvents, heavy metals and pesticides, diabetes, hypertension and obesity during midlife, tobacco use, depression, lack of cognitive engagement, or limited educational achievement (*5, 8, 9*).

To date, genome-wide association studies (GWASs) have identified over 80 loci linked with a higher sAD risk, with corresponding genes involved in several cellular processes, including endocytosis, cholesterol metabolism and immune response (*10–12*). Each of these loci contributes to an increase in risk, although the exact mechanisms by which they influence disease development remain often unclear. Thus, while polygenic risk scores can aggregate the small effects of multiple risk alleles (*12, 13*), creating disease models that accurately reflect the cumulative genetic risk remains difficult (*14–16*). Moreover, genetic factors alone do not fully explain the sAD onset and progression.

For this reason, human cell models derived from human induced pluripotent stem cells (hiPSCs), either derived from sAD patients or engineered to incorporate genetic risk variants, often exhibit only partial phenotypes of the disease. Increased Aβ40 secretion, elevated phosphorylated tau, and increased active GSK-3β were recently observed in neurons from a sAD patient (*16*). However, a different study found no elevation in secreted Aβ40 or Aβ42 in neurons from two sAD patients, although intracellular Aβ fragments were increased in one case (*17*). Similarly, no significant changes were observed in secreted Aβ levels or the Aβ42/Aβ40 ratio in sAD-derived neurons from three sAD patients (*18*). Other reports showed increased extracellular Aβ42 levels in sAD APOE-ε3/ε4 neurons, as well as increase in phosphorylated tau proteins and upregulation of GSK-3β (*19*). A recent study showed that sAD neurons with deficient function of REST (RE1-silencing transcription factor) demonstrated accelerated neural differentiation, reduced progenitor cell renewal and nuclear lamina disruption, which are phenotypes of early AD stages (*20*). CRISPR-engineered hiPSC models incorporating GWAS variants have provided additional insights. Knockout (KO) neurons for Bridging Integrator 1 (*BIN1*), showed disrupted calcium homeostasis and altered neuronal firing pattern (*21*). Engineered APOE-ε4 neurons displayed increased synapse numbers and Aβ42 secretion (*22*). A KO neuronal model of the polycomb complex protein BMI1 yielded one of the few cellular models reported literature for sAD showing accumulation of both Aβ42 and p-tau deposits (*23*). Moreover, when KO was performed in aged neurons *in vitro* (60 days), a decrease in both PSD-95 and synaptophysin levels could also be observed, suggesting a loss of synaptic integrity.

Overall, current sAD models generally do not to consistently reproduce strong phenotypes of the AD pathology, including robust Aβ and tau aggregation and pronounced structural or functional degeneration of synapses. Stem cell-derived models that exhibit aggregation of Aβ, for instance, need to rely on complex three-dimensional (3D) culture formats and the overexpression of several mutations involved in the amyloidogenic processing of APP (*14–16, 22, 24–28*). Aβ aggregates have been detected in cortical neurons differentiated from Down syndrome hiPSCs after 40 to 90 days in culture (*25*). A 3D neuronal model overexpressing *APP^KM670/671NL^*, *APP^V717I^* and *PSEN^ΔE9^* was able to produce Aβ plaques and neurofibrillary tangles after 50 to 80 days in culture (*29*). The overexpression of two copies of the *APP*, *PSEN^M136I^* and *PSEN^A264E^* generated Aβ aggregates in a hiPSC 3D neuronal model after 90 days of culture (*30*). Nonetheless, the phenotypes observed are often milder in 2D hiPSC-fAD neuronal cultures. Cortical neurons with the Swedish mutation in *APP* (*APP^Swe^*) showed a consistent rise in soluble Aβ monomer secretion, together with an increase in the Aβ42/Aβ40 ratio after 11 days *in vitro*, alongside with neuronal hyperexcitability at later stages of maturation. However, Aβ aggregation, tau hyperphosphorylation and concomitant effects on neuronal health are absent in this model (*31*).

Overall, the development of robust and comprehensive AD cellular models remains a significant challenge. As a common feature, current models often focus on genetic factors, rather than on the role of environmental triggers in AD pathogenesis. This limitation constrains the accuracy and translational potential of these systems, highlighting the need for more holistic approaches that capture the complex spectrum of AD pathological processes.

In this paper, we take into account both the genetic and non-genetic factors, which differ across the sAD patient landscape, that induce early functional changes leading to gene expression alterations. These alterations drive cells into specific disease states, which can be characterized by distinct transcriptomic fingerprints. Hence, the complexity of sAD may be described at the cellular level by defining early disease states characteristic of different patient cohorts with varying clinical trajectories. Despite divergent early states, these pathways may converge to manifest core late-stage disease hallmarks, including Aβ plaques, neurofibrillary tangles from hyperphosphorylated tau and chronic neuroinflammation (*7, 32, 33*). Defining the molecular identity of sAD cellular states, however, presents significant challenges. The brain inaccessibility in living patients restricts our ability to analyse early causative gene expression patterns directly. To address this problem, we analyse gene expression perturbations that may contribute to early disease onset and progression using transcriptomic data derived from early and late AD patients.

By screening dysregulated genes for their effects on sAD phenotypes, we identified *FBXO2*, a gene involved in APP degradation (*34*), from a transcriptomic signature derived from a ROSMAP cohort of AD patients (*35*). Downregulation of *FBXO2* in hiPSC-derived cortical neurons recapitulates Aβ aggregation, tau hyperphosphorylation, and structural and functional neuronal network impairment. These results indicate that the strategy adopted here can identify a set of early disease drivers and generate panels of preclinical models that capture the molecular complexity of the wide spectrum of sAD patients. The availability of such models is essential to inform on the early molecular aspects of the disease, provide insights into patient stratification, and aid in the identification of novel therapeutic targets.

## Results

### Analysis of early transcriptomic drivers of sporadic Alzheimer’s disease

We retrieved single-nucleus RNA-seq (snRNA-seq) data from a transcriptomic study performed on ROSMAP patient cohorts (*35*). The data were derived from the prefrontal cortex of 48 patients, comprising 24 healthy donors, 15 individuals with early AD and 9 individuals with late AD. These groups were classified based on cognitive function, amyloid and neurofibrillary tangle burden (*35*). The cell nuclei were categorised into different cell types and UMAP was employed to cluster these types within a 2D space, although no distinct patterning associated with AD progression was observed. Therefore, to enhance the granularity of the data, each cell type was further divided into sub-cell types using Louvain graph-clustering. Subsequently, the Pathifier method was utilized to assess how AD disrupts biological pathways within each sub-cell type by comparing them to healthy donors and generating disruption scores. With this method, the pathways that were perturbed in early AD stages were identified (see Methods). Among all the pathways identified, three were selected for further validation in this study, given its reported relevance in sporadic AD forms. These pathways were: (i) the synaptic vesicle cycle (*36*), (ii) the endo-lysosomal system (*37*), and (iii) the ubiquitin-mediated proteolysis (*38*). The specific genes within these disrupted pathways that contributed to a major extent to the perturbations were then analysed, pinpointing them as the potential early drivers of the pathology. The identified gene set for astrocytes, excitatory and inhibitory neurons is reported in **Figure 1A**. In this study, we focused on the validation of gene sets belonging to excitatory glutamatergic neurons, given that they were predicted to be one of the earliest affected cell types in AD (*39*). Additional genes were included on top of the bioinformatically-predicted list, such as the transcription factors EGR3 (*40*) or USF2 (*41*), given they reported role in controlling the expression levels of other genes present in our original list (e.g. SORCS3 or SNAP25 for EGR3 and lysosomal ATPase subunits for USF2).

**Figure 1.**
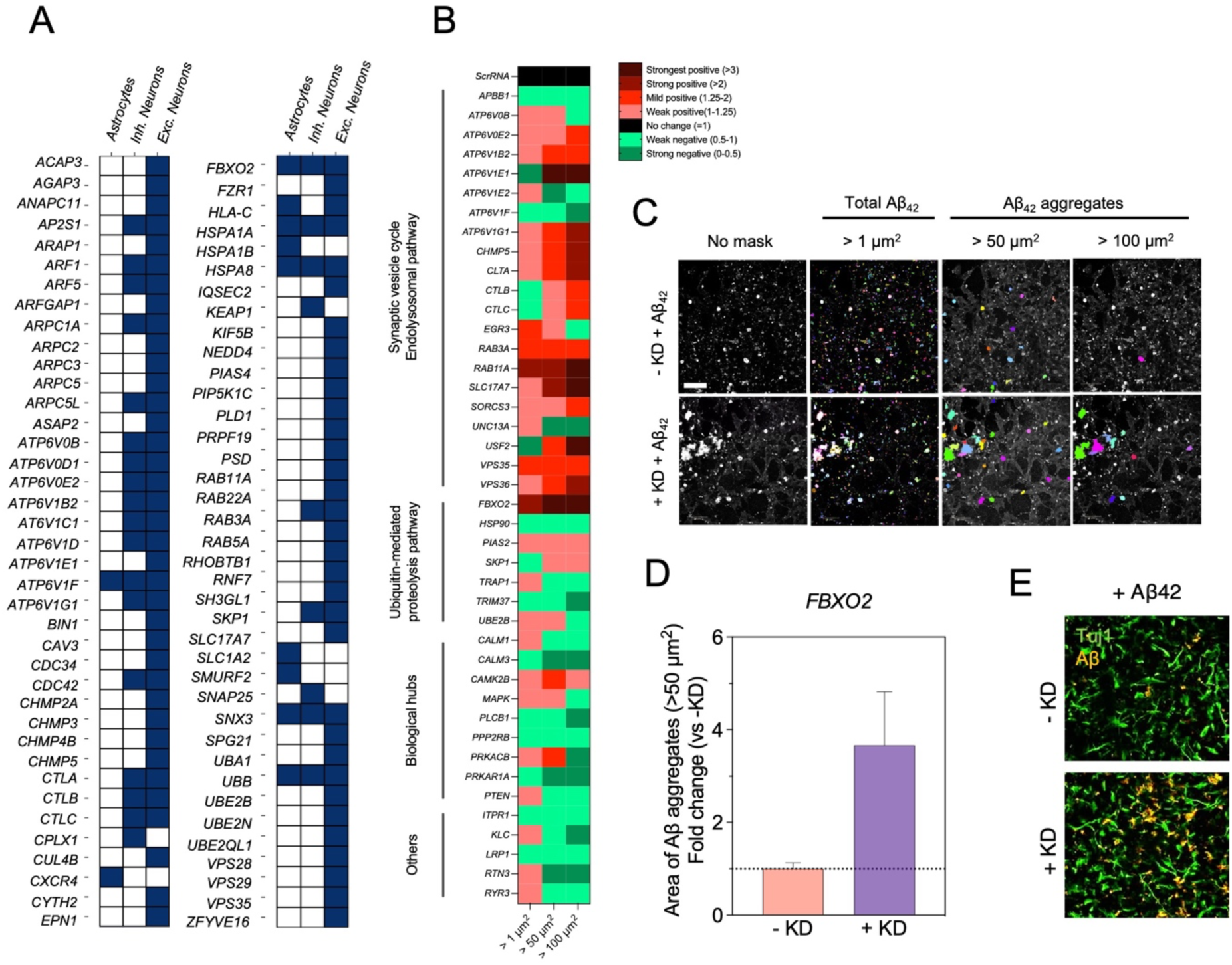
Screening in SH-SY5Y cells identifies genes whose downregulation enhances Aβ aggregation. **(A)** List of the 42 genes differentially expressed in early AD analysed in this study. **(B)** Test of these genes in SH-SY5Y cells to assess whether, upon downregulation, they increased the presence Aβ aggregates of different sizes (>1 µm^2^, >50 µm^2^ and >100 µm^2^). Screening results are represented in the form of a heat map, depicting the fold change in the total area occupied by each Aβ population relative to cells treated with scrambled RNA (or -KD). **(C)** Representative pictures of different masks settings generated with the Harmony software to select and quantify the extent of Aβ species with varying areas. **(D)** *FBXO2* downregulation resulted in over 3-fold increase in Aβ aggregates larger than 50 µm². **(E)** Representative pictures of the Aβ aggregates after *FBXO2* downregulation. Scale bar = 100 µm. n = 3 technical replicates.

### Identification of genes whose downregulation enhances Aβ aggregation in neuroblastoma cells

A total of 42 candidate genes (**Figure 1B** and Methods) were screened in wild-type human neuroblastoma (SH-SY5Y) cells by downregulation through RNA interference (RNAi). Upon knockdown (KD), monomeric Aβ was administered to mimic the imbalance between the generation and degradation pathways of Aβ and to trigger its deposition, as occurs in early AD stages (*7*). We then determined the effects of the downregulation of each candidate gene on Aβ aggregation and distribution (**Figure 1C**) by quantifying the total area of Aβ aggregates of three different sizes (>1 µm^2^, >50 µm^2^ and >100 µm^2^). Gene candidates that showed a mild-positive effect (fold change ranging from 1.25 to 2) in increasing the levels of large Aβ aggregates (>50 or >100 µm^2^) as compared to controls were selected for further validation. Following these criteria, we thus identified 16 genes whose downregulation increased Aβ aggregation levels in this system (**Table S1**). Most of these genes are involved in endocytosis, intracellular trafficking, and exocytosis. Key among these are numerous ATPase subunits found in the lysosomal membrane and transcription factors that control their expression, such as USF2 (*41*). These proteins play key roles in lysosomal compartment acidification for effective protein degradation. Alterations in the pH of endosomal compartments are linked to changes in APP processing and Aβ degradation, which may explain the observed increase in Aβ levels (*42*). Similarly, proteins that govern the fusion and intracellular transit of endosomes, including RAB3A, RAB11A, VPS36, and CHMP5 also increased Aβ aggregation levels after knockdown. APP, Aβ, and enzymes associated with APP amyloidogenic metabolism, like β-secretase, are cargoes of these organelles. Hence, disruptions in endosome generation, fusion, and degradation cycles might affect Aβ generation, intracellular retention time and degradation, increasing its levels (*43–45*). Among the shortlisted genes associated with the ubiquitin proteasome system, *FBXO2* downregulation demonstrated a strong effect in enhancing Aβ aggregation (**Figure 1D,E**). FBXO2 is a component of the SCF ubiquitin protein ligase complex that targets APP (*34*), although it is still unknown whether FBXO2 is directly involved in Aβ degradation processes. Our results indicate that *FBXO2* downregulation increases Aβ levels and its aggregation state.

### Validation of short-listed genes in hiPSC-derived glutamatergic neurons

To further validate the shortlist of gene candidates obtained in the SH-SY5Y screening, we chose four genes with different ability in enhancing Aβ aggregation and we tested them in hiPSC-derived glutamatergic neurons, a more physiologically relevant system.

For this purpose, a hiPSC clone (BIONi010-C-38) carrying the Swedish mutation on the *APP* gene (KM670/671NL, referred as APP^Swe^) together with the wild-type hiPSC HPSI0114i-KOLF2 (referred as APP^WT^) were banked and differentiated to cortical neuronal progenitor cells (NPCs). Human cortical NPCs from the two lines (BIONi010-C-38, HPSI0114i-KOLF2) were differentiated to mature glutamatergic cortical neurons and subjected to the respective quality controls (**Figures S1A,B**). NPC batches with high expression of the transcription factors EMX1 (>70%) and DLX5 (>40%), and low expression of NKX2.1 (<20%), were identified as cortical progenitors and utilised to generate glutamatergic neurons of the top layers of the cerebral cortex (*46*) positive for TBR1 (a cortical neuron-specific marker) (*47–49*), VGLUT1/2 (a glutamatergic neuron-specific marker), and negative for VGAT (a GABAergic neuron-specific marker) (*50*) (**Figures S1B,C** and **S2**). NPC-derived neurons from both phenotypes showed enrichment both in pre-synaptic and post-synaptic markers (synapsin-1 and PSD-95) (**Figure S2**) and demonstrated measurable electrophysiological activity (see below).

For gene candidate validation, a strategy based on periodic siRNA delivery at early stages of the neuronal maturation was optimised and tailored for the two lines used in this study (**Figure 2A**). Concretely, the delivery of the siRNAs was performed every three days, from DIV4 to DIV11. We then explored whether reducing their expression at early stages of the terminal differentiation of fAD and wild-type human glutamatergic neurons results in: (i) Aβ aggregation, (ii) tau hyperphosphorylation, and (iii) a functional neurodegenerative phenotype.

**Figure 2.**
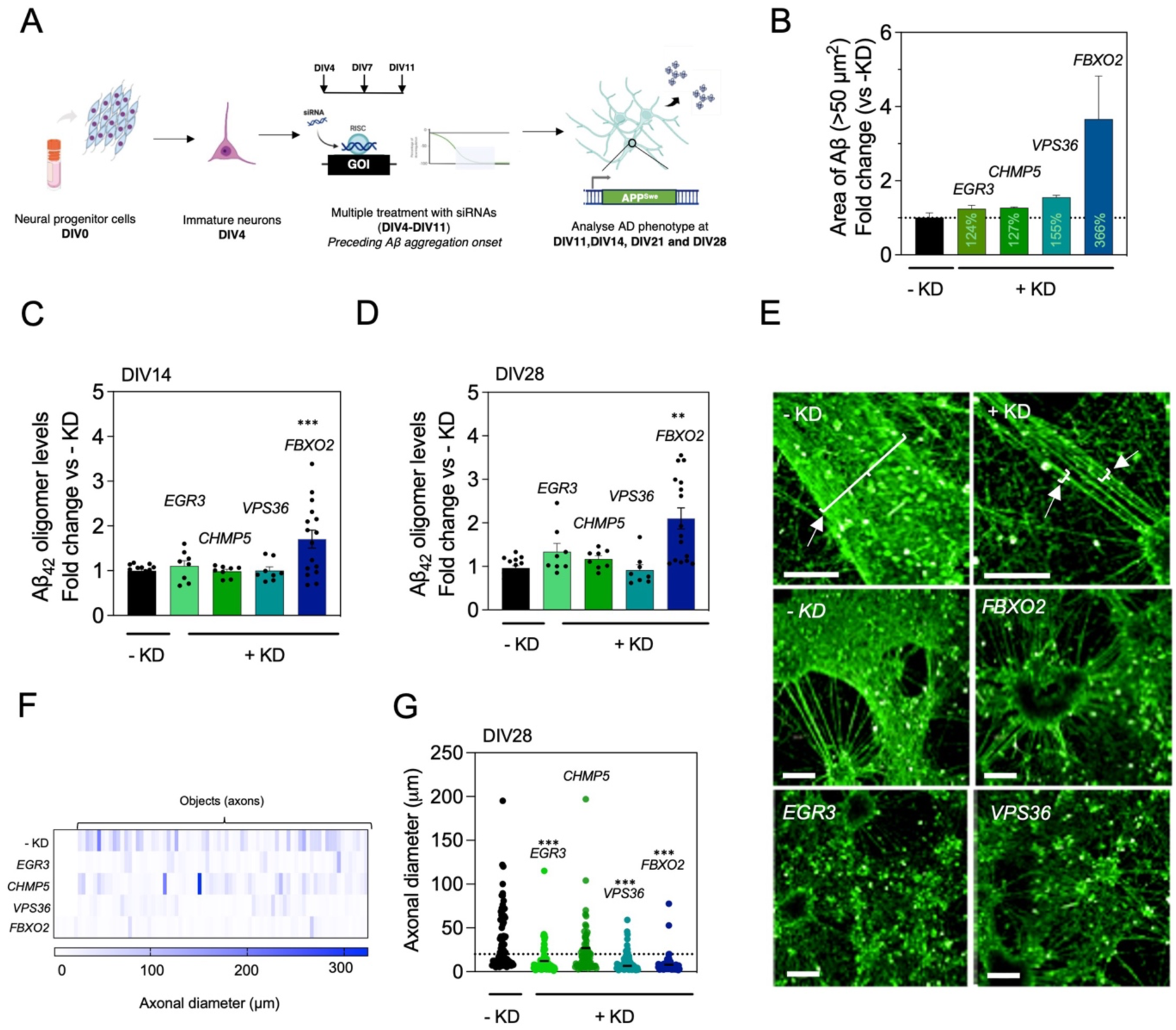
Screening of four short-listed genes in APP^Swe^ cortical neurons. **(A)** Strategy for downregulating candidate genes in hiPSC-derived APP^Swe^ neurons. Cortical neural progenitor cells (NPCs) were cultured and matured for 4 d to generate young neurons. Three siRNA administrations targeting each gene of interest (*EGR3*, *CHMP5*, *VPS36* and *FBXO2*) were conducted every 72 h (DIV4, DIV7 and DIV11). AD-relevant phenotypes were assessed at time points proximal to the siRNA intervention (DIV14) and at later stages of neuronal maturation (DIV28). **(B)** Fold change of the total area of Aβ aggregates larger than 50 µm^2^ of SH-SY5Y cells treated with 4 µM of monomeric Aβ42 subjected to a treatment with 10 nM siRNAs against *EGR3*, *CHMP5*, *VPS36* and *FBXO2*, calculated over the cells treated with 10 nM of scrambled RNAs (-KD). The percentage increase in Aβ aggregate area is depicted in each bar. (**C,D**) Fold change in secreted oligomer levels by APP^Swe^ neurons at 14 d (C) and 28 d (D) of maturation following the downregulation of *EGR3*, *CHMP5*, *VPS36* and *FBXO2*, calculated relative to the oligomer levels produced by non-KD APP^Swe^ neurons. *FBXO2* downregulation is the only condition that significantly increases Aβ oligomer production levels (n = 8-16). Absorbance values from oligomer detection were normalised by the mean cell density calculated from the wells where the supernatants were collected, calculated by measuring the total axonal (Tuj1) area. **(E-G)** Axonal diameter of DIV28 APP^Swe^ cortical neurons subjected to the downregulation of each screened gene -KD neurons were treated with scrambled RNA. Axons were stained with Tuj1 for assessment of their diameter (E). 75-100 axons were measured per condition. The diameter of each individual axon or axonal bundle is depicted in panels F and G. Notably, the downregulation of *EGR3*, *VPS36* and *FBXO2* triggered a statistically significant decrease in the axonal diameter. Statistical significance of ELISA and immunocytochemistry quantifications is indicated as: **p<0.01, ***p<0.001, determined by a one-way ANOVA test, further corrected for multiple comparisons using Bonferroni’s test.

The time window for the gene silencing was chosen in order to precede the detection peak of both Aβ and phospho-tau puncta in neurites (**Figures S2A** and **S4A**) both APP^Swe^ and APP^WT^ neurons expressed synaptic, axonal, and dendritic markers (e.g., Syn1, PSD-95, Tuj1, MAP2) and lacked NPC markers (e.g., Nestin, EMX1, PAX6).

### Screening of short-listed genes in APP^Swe^ hiPSC-derived neurons

Our screening approach was designed to evaluate the impact of early downregulation of each candidate gene in both fAD and WT hiPSC lines. This dual-line approach serves two distinct purposes. By downregulating candidate genes in fAD cells, with a strong genetic predisposition to AD, we aim to determine whether this early intervention exacerbates the disease phenotypes under investigation. If such exacerbation occurs, we posit that the early downregulation creates a molecular state of increased vulnerability to AD pathology. Conversely, applying the same early downregulation to WT “healthy” cells, lacking known genetic risk factors for AD, allows us to assess whether this induced vulnerable molecular state is sufficient to trigger disease-relevant cascades or phenotypes in an otherwise non-predisposed cellular system.

Therefore, following the procedure described above, we selected four genes to assess the effect of their KD in fAD APP^Swe^ hiPSC-derived neurons. The four selected genes were: (a) one longlisted gene (*FBXO2*) that scored high in the SH-SY5Y pre-screening, involved in UMP; (b) two longlisted genes (*VPS36* and *CHMP5*) that showed a mild effect on the increase of Aβ aggregation levels in the SH-SY5Y screening, involved in SVC and ELS, and (c) one gene (*EGR3*) that encodes a transcription factor that regulates the expression of numerous other candidates (*SORCS3*, *SNAP25*), reported to be downregulated in early AD, but that did not pass the cut-off of our SH-SY5Y screening assay (*40*) (**Figure 2B**). We examined two phenotypes, the secretion of Aβ oligomers and the alteration of the neuronal network structure. At early time points (DIV14), we detected higher levels of Aβ oligomers in the supernatant of neurons subjected to *FBXO2* downregulation, while the rest of the candidates showed no effects (**Figure 2C**). At later time points (DIV28), this increase prevailed in the *FBXO2* KD neurons, and a mild effect began to be observed for the *EGR3* and *CHMP5* downregulation conditions (**Figure 2D**). Alongside, we stained our neurons with Tuj1, an axon-specific marker, to measure the complexity of their axonal projections at DIV28. Without any stressors, APP^Swe^ cortical neurons clustered and formed notably thick axonal bundles upon maturation in the applied culture conditions. Some of these bundles even exceeded 50-100 µm in diameter (**Figure 2E**). This well-developed axonal structure was also evident in neurons that underwent treatment with scramble (labelled as -KD) or *CHMP5* siRNA duplexes (**Figure 2F,G**). In contrast, neurons subjected to *EGR3*, *VPS36*, and *FBXO2* downregulation showed thinner projections (**Figure 2F,G**), a phenotype that might relate to a potential alteration in electrophysiological function. Given this evidence, a more detailed analysis of the pathophysiological implications of *FBXO2* downregulation was performed in the APP^Swe^ neurons.

### *FBXO2* downregulation in APP^Swe^ cortical neurons triggers Aβ aggregation, tau hyperphosphorylation and functional network degeneration

We investigated the effects of an early downregulation of *FBXO2* in APP^Swe^ mutant human cortical neurons focusing on Aβ oligomer secretion, cell-associated Aβ, and phosphorylated tau levels, given the following considerations. The APP processing occurs at cell membranes, leading to the secretion of Aβ into the extracellular space. The accumulation of Aβ monomers in the supernatant promotes the formation of Aβ oligomers, which are considered among the most toxic species in AD (*7*). In addition, APP can also be endocytosed. In endosomes, APP undergoes amyloidogenic processing and generates Aβ that tends to accumulate intracellularly (*51, 52*), primarily in neurites.

The downregulation of the top candidate gene *FBXO2* was assessed by reverse transcription quantitative polymerase chain reaction (qPCR) (**Figure 3A**). Samples were taken 72 h after each siRNA shot. *FBXO2* mRNA levels were depleted by over 85% after two siRNA shots (DIV11). In the absence of any perturbation, the levels of secreted Aβ oligomers remained stable during APP^Swe^ neuronal maturation (**Figure 3B**). Conversely, *FBXO2* downregulation caused over a 2-fold increase in Aβ oligomerisation (**Figure 3C**) at all explored time points (from DIV11 to DIV28), which progressively raised upon neuronal ageing as compared to ScrRNA treated neurons (-KD). While the secretion levels of Aβ oligomers did not change in basal conditions, APP^Swe^ neurons exhibited a progressive accumulation of cell-associated Aβ levels over time. Aβ-derived signal, detected as bright spots primarily localized in neuronal axons, showed a peak at 14-days *in vitro*. Presumably, due to this elevated baseline level of intracellular Aβ in the mutant line, only a mild increase was observed in the formation of Aβ clusters in neuronal axonal projections both at DIV14 and DIV28 (**Figure 3D**). Similarly, APP^Swe^ neurons demonstrated a progressive increase in phosphorylated tau levels as they matured (**Figure S2A**). These levels began to be detected at early stages (DIV7) and seemed to reach a plateau by DIV14-21. We observed more pronounced tau phosphorylation at Ser396 following *FBXO2* knockdown both at DIV14 and DIV28 (**Figure 3E,F**). Notably, tau hyperphosphorylation at Ser396 has been reported in neurofibrillary tangles both at early and late stages of the pathology in human brain samples (*53*).

**Figure 3.**
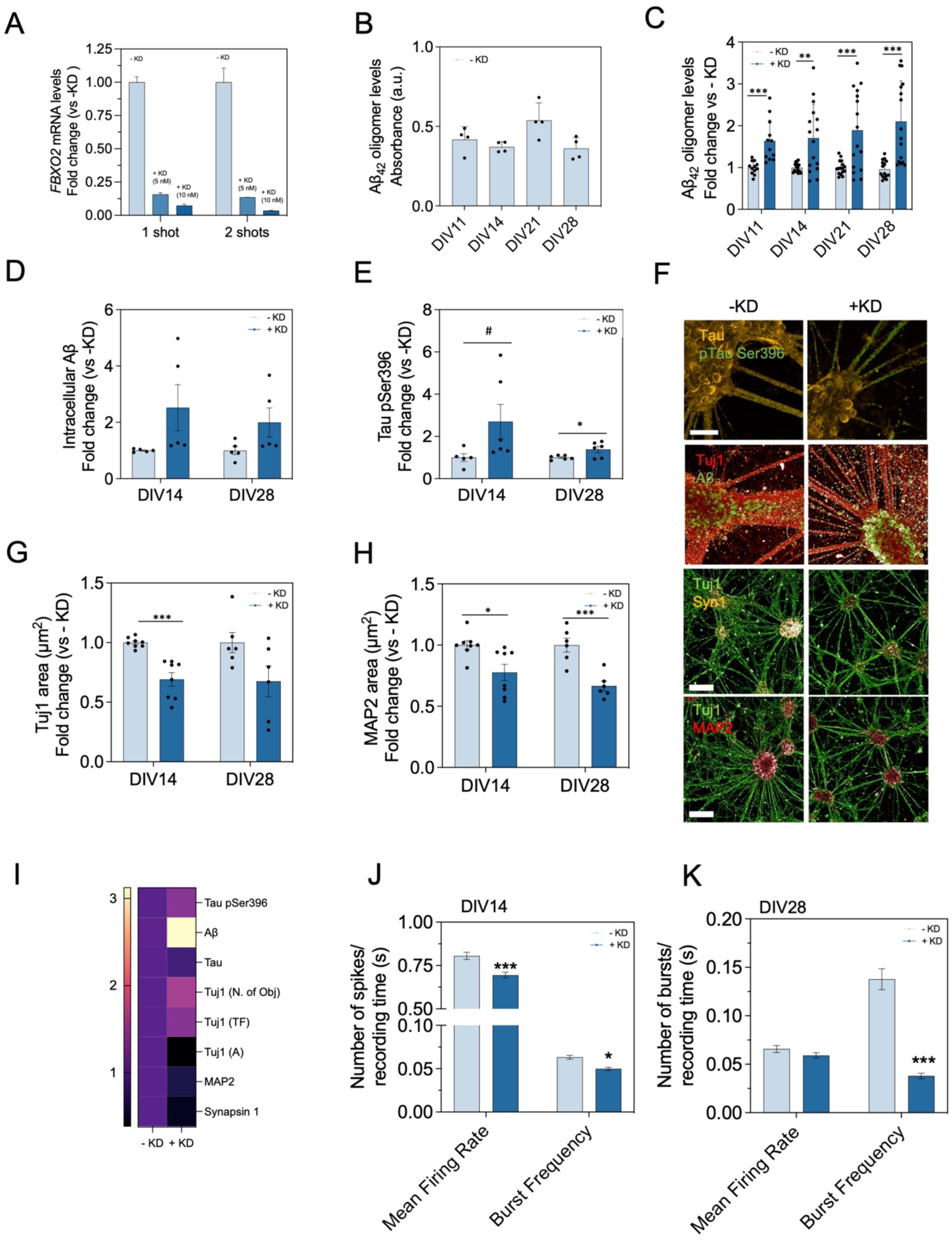
*FBXO2* downregulation worsens AD phenotypes in APP^Swe^ cortical neurons. **(A)** Validation of *FBXO2* downregulation. APP^Swe^ cortical neurons were subjected to a single (at DIV4) or double (at DIV4 and DIV7) siRNA treatment. Two different concentrations of siRNAs were used (5 and 10 nM). RNA samples were collected 72 h after each treatment. A single treatment was sufficient to downregulate *FBXO2* expression by over 80%. The ΔΔCt method was used for FBXO2 relative expression quantification compared to ScrRNA treated cells (-KD) (n = 3). **(B)** Homotypic 6E10-6E10 ELISA results illustrating Aβ oligomer levels detected in the supernatant of APP^Swe^ neurons at DIV11, DIV14, DIV21 and DIV28 without *FBXO2* downregulation (-KD), showcasing steady oligomer secretion levels over time. **(C)** Fold change in secreted Aβ oligomer levels by APP^Swe^ neurons at DIV11, DIV14, DIV21 and DIV28 following *FBXO2* downregulation, calculated relative to the oligomer levels produced by non-KD APP^Swe^ neurons at each time point. *FBXO2* downregulation increases Aβ oligomer production levels (N = 2, n = 12-16). The cell density was estimated from the wells where supernatants were collected by calculating the total Tuj1 area. These values were used to normalise oligomer absorbance values before fold changes were calculated. **(D,E)** Fold change of intracellular Aβ (D) and phosphorylated tau at Ser396 (E) levels located in the axonal projections of APP^Swe^ neurons at DIV14 and DIV28 after an early treatment with ScrRNA (-KD) or *FBXO2* siRNAs (+KD) (N = 2). Results indicate that *FBXO2* downregulation increases intracellular Aβ and phosphorylated tau levels. **(F)** Representative images of Aβ and pSer396 spots in APP^Swe^ neurons with or without *FBXO2* downregulation at DIV21. Scale bar = 50 µm for Aβ and phosphorylated tau panels. **(G-H)** Fold change decrease of the axonal (Tuj1, G) and dendritic (MAP2, H) area of APP^Swe^ neurons at DIV14 and DIV28 in the absence (-KD) or presence (+KD) of an early downregulation of *FBXO2*. Representative images are shown in panel (F) (N=2). Scale bar = 100 µm for Tuj1 and MAP2 panels. (**I**) Heat map of the fold change increase in AD-relevant protein aggregation markers (Aβ, pSer396 tau), coupled with a decrease in key markers of the neuronal network structure and function (MAP2, Tuj1, synapsin1, tau area) at DIV21 (n = 3). Increase in Tuj1 objects and Tuj1 fluorescence were indicators of axonal fragmentation. (**J,K**) Mean firing rate, and burst frequency measured as the number of individual spikes or high-frequency bursts of APP^Swe^ neurons with or without early *FBXO2* downregulation cultured in MEA plates and subjected to 5 minutes of recording of their electrical activity at DIV14 (J) and DIV28 (K). Statistical significance is denoted as *p<0.05, ***p<0.001, determined through a two-way ANOVA test, further corrected for multiple comparisons using Tukey’s test.

We then explored whether the presence of these Aβ and tau pathological traits triggered a loss of the structural neural network integrity and the neuronal electrophysiological function. As reported above, the APP^Swe^ neurons cluster and form thick axonal shafts (**Figure S5**). This phenomenon results in a reduction of the total axonal extension (referred to as the Tuj1 area) over time (**Figure S2**). Therefore, *FBXO2* downregulation led to a decrease in the total axonal area as compared to the control (-KD) exclusively at early stages of the APP^Swe^ neuronal maturation (DIV14), when the axonal network is still less compacted. However, at later time points, when the basal neuronal clustering is more pronounced, the decrease in the axonal area is no longer statistically significant (**Figure 3G**). Nonetheless, the downregulation of *FBXO2* led to axonal alterations, causing them to become thinner at DIV28 (**Figure 2**). This thinning of the axons was accompanied by a reduction in the size of the cellular clusters, where neuronal cell bodies and dendrites, rich in MAP2, gather. As a result, a reduced extension of MAP2^+^ dendritic projections were observed, a phenotype that was sustained through neuronal aging (from DIV14 to DIV28) (**Figure 3H**). Furthermore, the decrease in Tuj1^+^ area was coupled with an increase in the number of Tuj1 objects and Tuj1-derived fluorescence intensity (**Figure 3I**) indicating the presence of a bigger number of fragmented axonal projections, at DIV21. This perturbation also correlated with lower levels of synapsin-1 (**Figure 3I**), indicating a potential alteration in the synaptic activity of these neurons, concomitant with higher detectable levels of both Aβ and phosphorylated tau.

We then recorded the electrical activity of APP^Swe^ neurons subjected to the downregulation of *FBXO2* (+KD) and their corresponding control (ScrRNA, -KD) using multielectrode array (MEA) plates (**Figure 3J,K**). If neurons are active, their electrical firing is recorded by the electrodes located on the plate surface. Two parameters were analysed. First, the mean firing rate (Hz), which estimates the ratio between the total number of electrical spikes recorded and the duration of the recording. Second, the burst frequency, which is calculated as the number of neuronal bursts during the duration of the recording. With maturation, neurons fire action potentials in brief, high-frequency bursts, which are key for the effective transmission of information. Notably, at early time points (≃ DIV14), *FBXO2* KD neurons showed an impairment of their synaptic function, shown by a lower mean firing rate and burst frequency (**Figure 3J**). This impairment in electrophysiological activity upon *FBXO2* downregulation was maintained at later time points (≃DIV28) (**Figure 3K**).

Taken together, these data demonstrate that *FBXO2* downregulation pushes the neurons towards a neurodegenerative cascade recapitulating primary AD hallmarks: Aβ oligomer formation and aggregation, tau hyperphosphorylation, and synaptic dysfunction.

### Effects of *FBXO2* downregulation in APP^WT^ cortical neurons

Next, to investigate whether *FBXO2* can be used to model sAD in wild-type neurons, without genetic predisposition to AD, we implemented the strategy outlined in **Figure 4A** on APP^WT^ neurons. In short, we downregulated *FBXO2* expression (**Figure 4B**) in early neuronal maturation stages (DIV4 to DIV11) and analysed AD phenotypes at times close to (DIV14) and distant from the downregulation window (DIV21 and DIV28).

**Figure 4.**
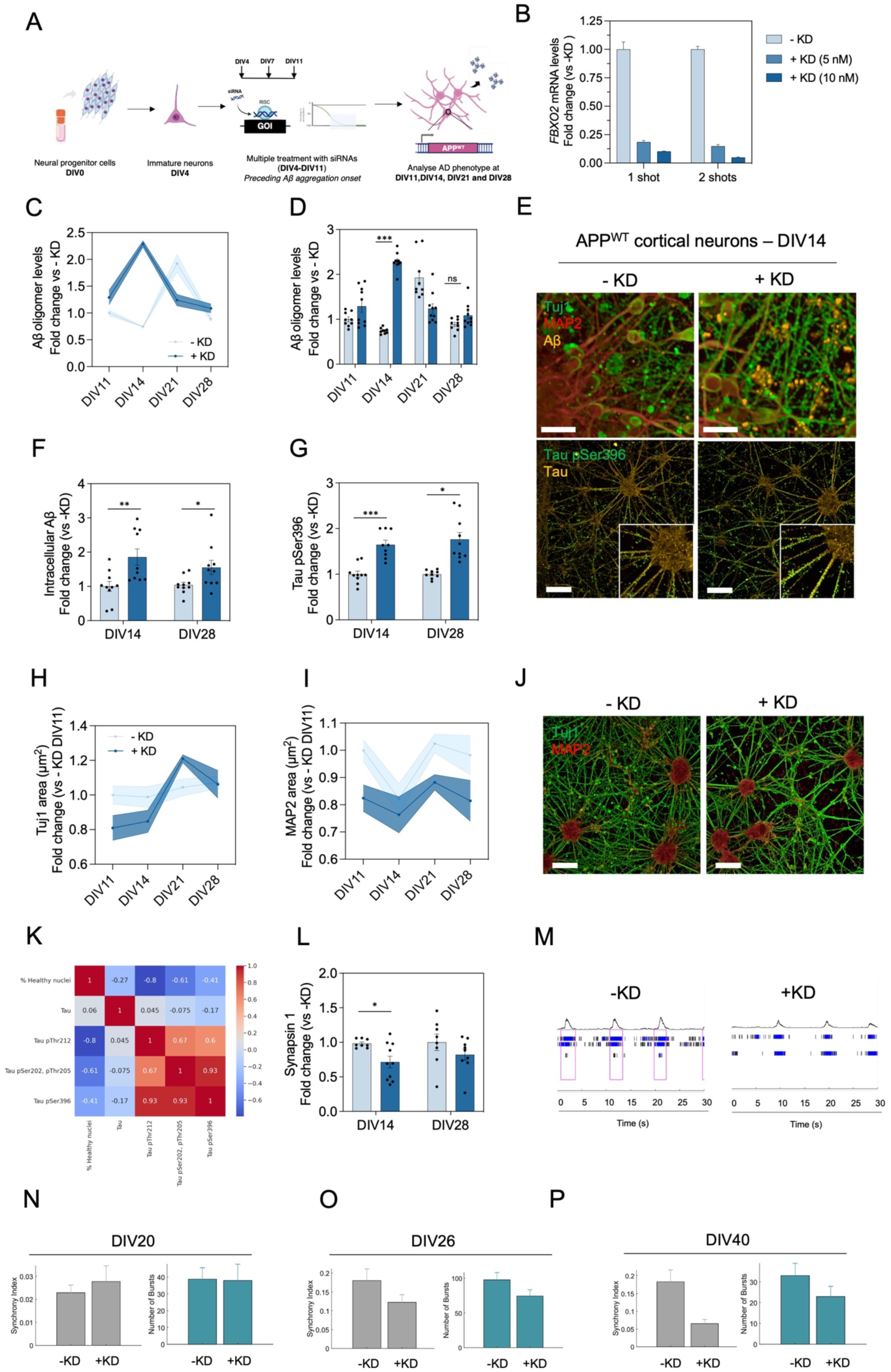
*FBXO2* downregulation induces AD phenotypes in APP^WT^ cortical neurons. **(A)** Strategy for downregulating candidate gene expression in hiPSC-derived wild-type neurons. Cortical neural progenitor cells (NPCs) were cultured and matured for 4 days to generate young neurons. Three siRNA administrations targeting the gene of interest (e.g., *FBXO2*) were conducted at 3-day intervals (DIV4, DIV7, DIV11). The AD phenotype was assessed at time points proximal to the siRNA intervention (DIV14) and at later stages of neuronal maturation (DIV21, DIV28), to investigate if gene downregulation propels the neuronal culture towards recapitulating Aβ and tau aggregation together with network degeneration. **(B)** Validation of *FBXO2* downregulation. As with the mutant line, APP^WT^ cortical neurons were subjected to a single (at DIV4) or double (at DIV4 and DIV7) treatment using siRNAs at two different concentrations (5 or 10 nM). 10 nM ScrRNA siRNAs were used as a negative control. Samples were collected 72 h after each treatment. Relative expression quantification for FBXO2 was calculated by the ΔΔCt method (n = 3). **(C-D)** Progression of Aβ oligomer levels detected in the supernatant of APP^WT^ neuronal cultures with or without *FBXO2* downregulation, by performing a homotypic 6E10-6E10 ELISA. Data are presented as the fold change calculated relative to the mean absorbance measured from the supernatant of APP^WT^ neurons without *FBXO2* downregulation (-KD) at DIV11. *FBXO2* downregulation induces an earlier and more pronounced peak of oligomer secretion as compared to control (N=2, n=10). **(E)** Representative images showing an increase in intracellular and membrane-associated Aβ and phosphorylated tau (Ser396) levels of APP^WT^ cortical neurons at DIV14 subjected to an early treatment with ScrRNA or *FBXO2* siRNAs. Scale bar = 50 µm. **(F,G)** Quantifications are shown as the fold change increase in the total count of either Aβ (F) or phosphorylated tau (G) spots located in axonal projections, normalised by the total Tuj1 area (N=3), at early (DIV14) and late (DIV28) time points. **(K)** Matrix reporting the negative correlations between the neuronal health (estimated by the percentage of healthy nuclei) and the levels of phosphorylated tau at different residues (Thr212, Ser202-Thr205, and Ser396) in *FBXO2* KD APP^WT^ neurons at DIV14. **(L)** Quantification of the fold change decrease in synapsin 1 positive area in FBXO2 KD APP^WT^ as compared to the control (-KD), statistically significant at early time points (DIV14) (N=3). **(M)** Raster plot from a MEA plate well depicting the electrical activity of APP^WT^ neurons with or without an early downregulation of *FBXO2* at DIV26. Each row represents electrical activity recorded from a single electrode. Individual asynchronous spikes are shown as black marks, whereas bursts recorded from a single electrode appear as blue marks. Synchronised bursts across multiple electrodes are highlighted in a pink square. The synchrony of neuronal bursts is lower upon *FBXO2* downregulation. **(N-P)** Quantification of both the synchrony index and the number of bursts from 32 independent MEA wells measured at DIV20 (N), DIV26 (O), and DIV40 (P) in -KD and +KD APP^WT^ showcasing a mild and late electrophysiological function impairment. Statistical significance of ELISA and immunocytochemistry quantifications is indicated as: ^ns^p>0.05, *p<0.05, **p<0.01, ***p<0.001, determined by a two-way ANOVA test, further corrected for multiple comparisons using Tukey’s test.

As with APP^Swe^ neurons, after rigorous quality control on NPCs (**Figure S3**) and glutamatergic neurons (**Figure S4**), we examined Aβ oligomer production, intracellular Aβ aggregation and tau hyperphosphorylation. To explore the temporal dynamics of Aβ aggregation upon APP^WT^ neuronal aging, we quantified Aβ oligomer secretion across different days *in vitro* (DIVs 11, 14, 21 and 28) (**Figure 4C,D**). Notably, we observed an early, pronounced peak of Aβ oligomer production in *FBXO2* KD neurons (at DIV14) preceding a the one for -KD neurons (at DIV21). Aβ oligomer levels detected in the supernatant diminished over time, with differences lost between KD and non-KD groups at DIV28 (**Figure 4D**). This acute, short-lasting effect observed in the WT was likely due to lack of any source of genetic alteration promoting abnormal cleavage of APP and increasing Aβ secretion to the extracellular space, as occurs with the Swedish variant on the *APP* gene. APP^WT^ neurons (-KD) did progressively accumulate intracellular Aβ aggregates in their cell bodies and axonal projections (**Figure S4**). These levels were significantly increased upon the downregulation of *FBXO2*, particularly at earlier time points (DIV14) (**Figure 4E,F**). However, the absolute levels of Aβ were lower than those observed in the APP^Swe^ neurons (**Figure S5**). Specifically, the Aβ levels in APP^Swe^ neurons were over 50% higher than in APP^WT^ neurons. On the other hand, the levels of tau phosphorylation at Ser396 also increased significantly upon *FBXO2* downregulation, similarly to what we observed in the mutant line (**Figure 4E,G**). These results suggest that, in the absence of a familiar AD mutation, *FBXO2* downregulation opens a window of increased vulnerability for the onset of Aβ aggregation and tau hyperphosphorylation.

We then examined the effects of *FBXO2* downregulation on neuronal health and function. The amyloid hypothesis proposes that tau hyperphosphorylation is a downstream event resulting from Aβ aggregation (*54*). Elevated tau phosphorylation levels increase its propensity to aggregate and trigger cell death (*55*).Therefore, we studied the correlation between the percentage of unhealthy nuclei in *FBXO2* KD neurons and tau phosphorylation levels at three sets of residues (Thr212, Ser202-Thr205, and Ser396) (**Figure 4K**). The unhealthy nuclei population was defined based on the morphology and packing density of the neuronal nuclei chromatin, which can be visualized through Hoechst staining. First, we observed a positive correlation between phosphorylation of all the examined residues. The total levels of tau did not correlate with an increase or decrease in phosphorylated tau levels, suggesting that this phosphorylation likely depends on kinase activation or phosphatase inhibition driven by *FBXO2* downregulation. On the other hand, we observed a negative correlation between tau phosphorylation and the percentage of healthy nuclei, with this relationship being more pronounced for pThr212. We note that previous studies already reported that tau phosphorylation solely at this residue is sufficient to enhance the propensity of tau for self-assembly and initiate neurodegenerative cascades (*56*), as observed in our assay.

Concerning the structural network, APP^WT^ neurons showed a different axonal and dendritic distribution pattern upon maturation as compared to their APP^Swe^ counterparts. APP^WT^ neurons used in this study exhibited less clustering and develop increasing axonal branching, which grows in extension and complexity over the neuronal maturation course. This growth results in a progressive increase in the Tuj1^+^ area from DIV4 to DIV21 (**Figure S4**). Contrarily, the levels of MAP2 did not change significantly across different time points (**Figure S4**). Upon *FBXO2* downregulation, we observed a clear reduction in the extension of axons (DIV11-14, **Figure 4H,J**), a phenotype that mitigated at later time points (DIV28). We also noticed a reduction in MAP2^+^ dendritic regions (**Figure 4I,J**), which persisted through neuronal maturation, as also observed in *FBXO2* KD APP^Swe^ neurons.

We next analysed the effects of *FBXO2* downregulation on the APP^WT^ synapsin-1 levels and neuronal electrophysiological activity. We observed a mild decrease in the synapsin-1-derived signal upon *FBXO2* downregulation, which was statistically significant at early stages (DIV14) and remained present at later time points (DIV28) (**Figure 4L**). Concomitantly, the neurons presented a mild disruption in their synchronised firing patterns, or bursts, as defined by the synchrony index (**Figures 4M-P**). This perturbation was only observed at later stages, starting from 26-days *in vitro* and becoming more noticeable at DIV40. Alternatively, the perturbation in the neuronal network function was detected at earlier maturation stages of the APP^Swe^ *FBXO2* KD neurons (DIV14). This finding underscores the increased susceptibility of this phenotype to disease progression compared to our wild-type (WT) system.

Overall, we consistently observed that reducing the levels of expression of *FBXO2* elicits AD-relevant cascades in both APP^Swe^ and APP^WT^ neurons. Our findings, together with previous evidence (*57*), demonstrate that *FBXO2* downregulation can be used to model sAD in neurons, by triggering Aβ oligomerisation, intracellular Aβ aggregation, tau aberrant phosphorylation and neuronal dysfunction.

### Effects of *FBXO2* downregulation in APP^WT^ cortical neurons in brain-mimicking cellular environments

To prolong the observed AD phenotypes upon *FBXO2* KD and increase the physiological relevance of our sAD model, we introduced a series of modifications into our protocol (**Figure 5A,B**). First, we shifted from performing 50% medium changes to a top-up medium change regime (**Figure S6**). This regime involved the progressive addition of fresh medium supplemented with differentiation factors, without any medium removal, to prevent the continuous depletion of a fraction of the soluble Aβ species produced during the maturation and ageing of the cortical neurons. The top-up regime started at DIV14 (72 h after the last siRNA shot) and was kept until 33 days *in vitro*. Second, we modelled the astrocytic contribution to neuronal survival, maturity, and function (*58*), mediated by secreted neurotrophins and factors, by treating *FBXO2* +KD and -KD neurons with conditioned medium from healthy human astrocytes (ACM), starting from DIV14 (**Figure 5A,B**). Neurons aged with a 50% medium change regime using neuronal maintenance medium (NM) were used as reference.

**Figure 5.**
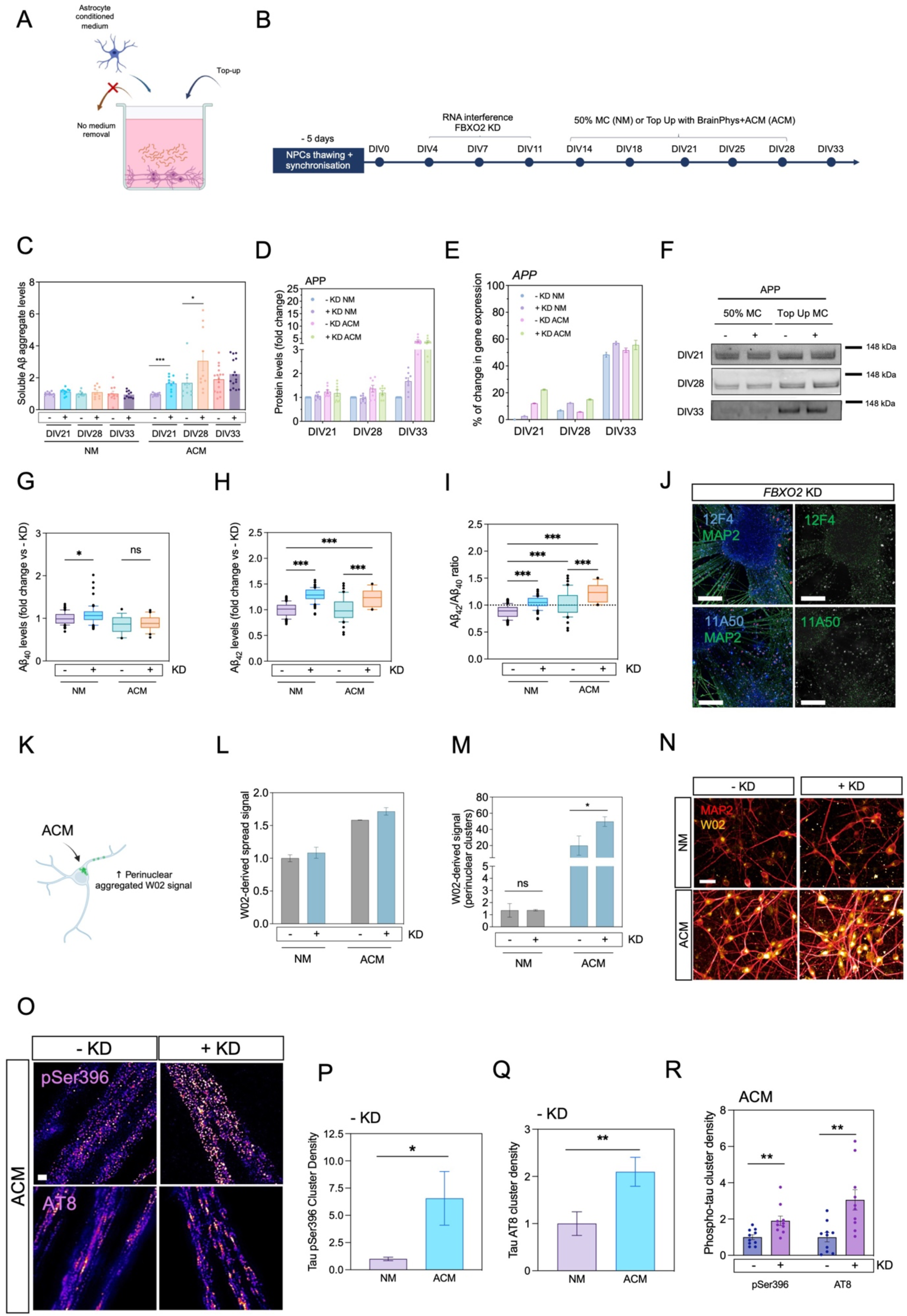
*FBXO2* downregulation induces characteristic AD phenotypes in APP^WT^ cortical neurons in AD-mimicking cellular environments. **(A,B)** Scheme of the experimental design. In brief, *FBXO2* is downregulated at early neuronal maturation stages (from DIV4 to DIV11). From DIV14, a top-up medium change is performed by adding a mixture of astrocyte conditioned medium (ACM) and neuronal maintenance medium. In parallel, neurons subjected to a 50% medium change regime with neuronal maintenance medium (NM) were cultured as reference. **(C)** Soluble Aβ aggregates quantified in the supernatant of NM and ACM-treated neurons at DIV21, DIV28 and DIV33 by a 6E10-6E10 homotypic ELISA (N = 2, n = 10). For each time point, statistical differences were calculated using a two-way ANOVA assay, comparing +KD with -KD for NM and ACM-treated cells, respectively, using Šídák’s multiple comparisons test (*p<0.033, ***p<0.001). **(D)** APP levels were measured by western blot from two independent biological replicates. The band intensity was analysed by ImageJ at least three times per replicate, to account for variability derived from image quantification. Intensity was normalised by the mean intensity of Ponceau-S per sample. **(E)** Percentage of change in *APP* mRNA levels (calculated via ΔΔCt values normalised by ACTB) in NM or ACM-treated neurons subjected to an early treatment with ScrRNA (-KD) or *FBXO2* siRNAs (+KD) over time, calculated vs -KD NM DIV21 neurons. **(F)** Blots showing APP levels over different time points and treatments. **(G-H)** Levels of Aβ40 (G) and Aβ42 (H) puncta, at DIV33, measured by immunocytochemistry using the 11A50B and 12F4 antibody, respectively, normalised by neuronal area. Data are represented as the fold change vs -KD NM treated neurons (n = 3, 27-36 independent images per technical replicate). **(I)** Aβ42/Aβ40 ratio for each condition, calculated with the normalised puncta levels for both peptides (n = 3). Statistical analyses for panels G-I were calculated using a two-way ANOVA, using Tukey’s test to correct for multiple comparisons test (*p<0.033). **(J)** Representative images of the levels and distribution of 12F4 and 11A50-positive puncta in ACM-treated neurons with *FBXO2* KD, at DIV33. Scale bar = 50 µm. Panels on the right show the cleaned masks (in green) for Aβ puncta, and the removal of bigger artifacts (in white) for quantification. **(K)** Diagram summarising that ACM-treated neurons demonstrate an increase in accumulation of perinuclear W02-positive signal of high intensity. **(L)** Quantification of the perinuclear W02 positive area independently of intensity at DIV33. Data are normalised by total MAP2 area. **(M)** Quantification of the total area occupied by high intensity W02-positive perinuclear clusters, normalised by the total MAP2 area, at DIV33. Statistical analyses as calculated using a two-way ANOVA, using Tukey’s test to correct for multiple comparisons (*p<0.033). **(N)** Representative images of high-intensity W02-positive perinuclear clusters in each treatment condition at DIV33. Scale bar = 20 µm. **(O)** STED images of phosphorylated tau at Ser396 and Ser202/Thr205 (AT8) of +KD and -KD ACM-treated neurons. Scale bar = 1 µm. **(P-Q)** Quantification of the puncta levels of phosphorylated tau Ser396 (P) or Ser202/Thr205 (Q), normalised by total MAP2 area, of DIV33 neurons with no early gene downregulation (-KD) cultured with NM or ACM protocols. Statistical differences were calculated with a Student’s t-test (* p<0.033, ** p<0.02). **(R)** Phosphorylated tau cluster density normalised by total tau-positive area. Each dot corresponds to the normalised phosphorylated tau levels calculated from an image (n = 10 images per condition). Statistical differences were calculated with a Student’s t-test (** p < 0.02).

*FBXO2* KD in WT neurons, following the original neuronal medium change regime protocol (NM), did not result in increased soluble Aβ oligomer production at later time points (DIV21-DIV33; **Figure 5C**), as also reported in **Figure 4C**. In contrast, ACM-treated neurons exhibited a time-dependent increase in Aβ oligomer generation, which was more pronounced upon *FBXO2* KD, both at DIV21 and DIV28 (**Figure 5C**). Notably, in both +KD and -KD conditions, ACM treatment led to increased APP protein levels as compared to NM-treated neurons, particularly at later time points (DIV33), assessed by western blot. Contrarily, ACM-treatment increased *APP* mRNA expression levels just at earlier time points (DIV21) but not at DIV28 or DIV33. These results suggest the components of ACM, after two-weeks of treatment, may alter APP degradation pathways in treated neurons, contributing to the higher Aβ oligomer levels observed at later stages (**Figures 5F-E, S7**).

Both NM and ACM-treated cells also displayed increased Aβ42 puncta in neurites and neuronal bodies following *FBXO2* KD (**Figures 5G-J**). Notably, the combination of ACM treatment and *FBXO2* KD resulted in a statistically significant increase in the Aβ42/Aβ40 ratio (**Figure 5I**). Moreover, we observed significant perinuclear accumulation of W02-positive signal, which was 50% higher in ACM-treated cells compared to NM-treated cells, accompanied by marked differences in signal intensity and morphology (**Figure 5K-N**). High-intensity W02 clusters were significantly elevated in ACM-treated cells, particularly with *FBXO2* downregulation, with fold changes exceeding 20 (**Figures 5M-N**). To note that, given that the W02 antibody binds to all forms of Aβ and APP, these perinuclear clusters may be a product of the contribution of all such protein species.

Additionally, ACM treatment also increased tau phosphorylation at residues Ser396 and Ser202/Thr205 compared to NM-treated cells (**Figures 5P,Q**). Using STED microscopy, we found that *FBXO2* downregulation in ACM-treated cells further elevated phosphorylated tau levels across all assessed residues (**Figures 5O-R**).

Taken together, these results underscore two primary findings. First, pathophysiological aspects of sAD, such as neuronal ageing (by extending the culture length), elevated Aβ retention (by applying a *top up* medium change regime) or the presence of astrocyte-derived factors help recapitulate characteristic features of the disease in human neuronal cell systems, such as Aβ or tau accumulation and aggregation. Second, even when employing more complex protocols that elicit stronger phenotypes in neuronal cells, the disease-worsening effects from the transient, early downregulation of *FBXO2*, persist.

## Discussion

Given the advances made in the development and application of RNA sequencing methods in the past decade, several studies have analysed the transcriptomic changes associated with sAD (*35, 59–63*). However, there is still a need for cellular systems to validate and integrate putative causative gene into sAD models. To address this problem, in this work we established a pipeline that translates predictions of early drivers of sAD into the generation of human cell models. With this pipeline, we validated the predictions using an AD SH-SY5Y model together with fAD and wild-type cortical glutamatergic neurons derived from hiPSCs. By using cellular systems with diverse genetic and molecular backgrounds, we aimed to enhance the robustness of the selected transcriptomic perturbations for model development, as their effects should promote AD phenotypes across multiple backgrounds.

The initial screening system was based on human neuroblastoma cells (SH-SY5Y). Although these cells are known to resemble human neurons, they exhibit low levels of endogenous Aβ production and aggregation, a key readout for gene validation in our study. To overcome this problem, we added exogenous Aβ monomers to promote basal Aβ aggregation and then assessed the impact of the gene knockdown on Aβ aggregate levels. Observed aggregates were likely to result from both exogenous and secreted Aβ, with the latter presumably exacerbated by the downregulation of specific candidate genes. Genes increasing Aβ aggregation in this system were selected for further validation. We note that the SH-SY5Y assay was used as a rapid system to select gene candidates for further analysis, and we did not use this assay to investigate the mechanisms by which each downregulated gene triggered aggregation in these cells.

We then selected the genes most effective in promoting Aβ aggregation upon downregulation in the SH-SY5Y assay, and assessed the effect of their knockdown in hiPSC-derived glutamatergic cortical neurons. Our goal was to use glutamatergic cortical neurons to: (i) mimic transcriptomic perturbations at early disease stages, preceding the manifestation of Aβ aggregation and tau hyperphosphorylation, and (ii) to assess the effects on these disease phenotypes induced by the knockdown of the gene candidates.

To design a workflow for this validation, we first defined an early window for gene expression perturbation, tailored to the two hiPSC lines used in this study. We then selected the earliest possible window during the terminal differentiation stage, where neurons already expressed synaptic, axonal, and dendritic markers (Syn1, PSD-95, Tuj1, and MAP2) and lacked NPC markers (nestin, EMX1, PAX6). Next, we chose a window preceding the basal peak levels of Aβ or phosphorylated tau puncta in the neurites. With these considerations, we periodically delivered siRNAs from DIV4 to DIV11, every three days, to silence gene expression for at least a week. After DIV11, we stopped siRNA delivery and allowed the neurons to age. We hypothesized that if a transcriptomic perturbation increases vulnerability for disease onset, the neurons are more likely to manifest AD-relevant phenotypes at later stages. To account for cytotoxic or nonspecific effects from siRNAs, we always included treatments with ScrRNA as a control.

Our validation began with a fAD line carrying the Swedish mutation in the APP gene (APP^KM670/671NL^ or APP^Swe^). APP^Swe^ cortical neurons exhibit a robust increase in soluble Aβ monomer secretion and an elevated Aβ42/Aβ40 ratio after 11 days *in vitro*, along with neuronal hyperexcitability at later maturation stages (*31*). However, this model does not readily recapitulate phenotypes related with more early disease stages, such as Aβ aggregation, tau hyperphosphorylation, and associated effects on neuronal health. In fact, most of the stem cell models for AD with robust Aβ aggregation and concomitant neurodegenerative phenotypes reported in literature are based on the overexpression of several mutational variants involved in the amyloidogenic processing of APP (*14–16, 22, 24–28*). Furthermore, such models are often not scalable for use as preclinical models for drug screening.

Given the predisposition of the APP^Swe^ neuronal model to mild disease phenotypes, we used it to determine if the downregulation of candidate genes could exacerbate these conditions. Under the reported culture conditions, the APP^Swe^ hiPSC clone did not show an increase in soluble Aβ oligomers over time, but it exhibited high clustering and thick axonal shafts upon maturation. Using these features as benchmarks, we evaluated the impact of early gene downregulation on two key parameters: (i) Aβ oligomer secretion in the supernatant, and (ii) the diameter of axonal projections. Among the genes tested, *FBXO2* knockdown resulted in both increased oligomerization and reduced axonal thickness. Notably, the effects on Aβ oligomer production in the supernatant persisted throughout the experiment (from DIV11 to DIV28). This was accompanied by a mild increase in neurite-associated Aβ puncta and a more pronounced increase in phosphorylated tau at residue Ser396, a target of GSK3β, which is strongly associated with AD pathology. Additionally, structural alterations in the axonal projections of *FBXO2* KD neurons correlated with decreased firing rate and burst frequency, tested in a different set of experiments. Taken together, these results show that the early downregulation of *FBXO2* exacerbates disease phenotypes across various experiments and readouts in APP^Swe^ cortical neurons.

We then tested the downregulation of *FBXO2* in wild-type glutamatergic cortical neurons to determine whether or not we could recapitulate disease-relevant hallmarks in the absence of a strong genetic predisposition for AD. Early *FBXO2* knockdown triggered an increase in oligomer secretion only at time points close to the siRNA intervention window (DIV11 and DIV14), with effects disappearing at later stages. Notably, we observed a significant increase in Aβ puncta in neurites and phosphorylated tau at both early and late time points. *FBXO2* KD also triggered a mild decrease in axonal and dendritic levels, lower expression of pre-synaptic markers such as Syn1 and impaired electrophysiological activity. Altogether, we demonstrated that *FBXO2* KD also increases the vulnerability of wild-type neurons to disease. We further modified the culture conditions of our *FBXO2* KD wild-type neurons to better mimic the composition of the brain extracellular space. For that, we applied a top-up medium change regime after 14-days *in vitro*, to prevent removal of soluble Aβ secreted by the cells, using a mixture of neuronal basal medium and human astrocyte conditioned medium. We observed that astrocyte conditioned medium was able to increase APP protein levels in treated neurons. By doing so, we promoted and prolonged the effects on Aβ oligomerization, intracellular Aβ aggregation, and tau phosphorylation up to 28 to 33 days *in vitro*, the latest time point explored in this study.

Overall, these results indicate that the human neuronal model of sAD that we developed by downregulating *FBXO2* in cortical neurons derived from hiPSCs replicates key molecular hallmarks of sAD, including Aβ aggregation, tau hyperphosphorylation, and functional network impairment. The E3 ubiquitin-ligase subunit FBXO2 has been shown to regulate the protein levels of three key AD targets: BACE1 (*64*), APP, and NMDA receptors (*34*). Some reports identified FBXO2 as a central multi-omic AD biomarker (*65*) and highlighted its potential as a therapeutic target (*66*). The identification of *FBXO2* as an important player in Aβ and tau pathology offers novel opportunities for targeted drug development, potentially leading to effective therapeutic interventions, and supporting the development of precision medicine approaches tailored to the heterogeneous nature of sAD.

In perspective, as the modulation of a single gene is unlikely to fully model a multifactorial disease, we anticipate that the approach that we described for *FBXO2* could be extended to incorporate other genes. In this way, it may be possible to create a panel of sAD neuronal models to capture the complex nature of the early molecular events leading to disease manifestations. Unlike fAD models, which rely on overexpression of causative genes, this approach reflects a broader spectrum of disease mechanisms driven by a combination of genetic and environmental factors, thus enhancing our understanding of the aetiology and progression of sAD.

Future research may focus on comparative studies with fAD models to delineate the distinct and overlapping pathways involved in sAD and fAD. Furthermore, integrating this model with other translational research tools, such as multiple patient-derived hiPSC lines, long-term stable knockdown methods using CRISPR/Cas9, mouse models and organ-on-a-chip technologies, could provide a more comprehensive understanding of the sAD pathology at the preclinical level, and offer more confidence that our observation in hiPSC-derived neuronal models could extend to the complex environment of the human brain.

In conclusion, the ability to capture the multi-factorial nature of sAD through human stem cell models through strategies of the type presented here will create a robust platform for preclinical studies of drug candidates, as well as opportunities development of personalized treatment strategies, tailored to individual genetic and environmental risk factors. As the field will continue to refine and validate these preclinical models, more tools will be built to enable the development of effective interventions that can alleviate the burden of AD and other neurodegenerative conditions.

## Materials and Methods

### Study design

The three major aims of this study were: (i) to identify early gene expression perturbations that can be drivers of sporadic Alzheimeŕs disease (sAD) from patient-derived transcriptomic data, (ii) to subject the predictions to three validation stages using human cellular systems to identify drivers of sAD pathology, and (iii) to use the early downregulation of the top hit (*FBXO2*) to trigger a set of sAD phenotypes in wild-type cortical neurons. To achieve these aims, we analyzed snRNA-seq data from the prefrontal cortex of 48 ROSMAP patients, categorized into healthy donors, early AD, and late AD (*35*), and identified cell types and sub-cell types using UMAP (*67*) and Louvain graph-clustering (*68*). Using the Pathifier method (*69*), we found disrupted pathways in specific sub-cell types in early versus late AD stages. From the list of predicted pathways, we focused on the synaptic vesicle cycle, endo-lysosomal system, and ubiquitin-mediated proteolysis, and pinpointed key genes driving these perturbations. We screened the effect of the resulting 42 predicted genes in SH-SY5Y cells. Using RNAi to downregulate gene expression, we then exogenously administered Aβ and assessed changes in Aβ aggregation levels by immunocytochemistry and high-content confocal microscopy. A portion of the genes were screened in APP^Swe^-derived cortical neurons. The selected candidate, *FBXO2*, was further validated in both APP^Swe^ and APP^WT^ neurons, derived from two different hiPSC lines respectively (BIONi010-C-38 and Kolf2-C1). A chemical differentiation protocol, as detailed below, was applied to convert hiPSCs into cortical NPCs. NPCs were subjected to quality controls prior their used for screening and validation (**Figures S1-S4**). It was of particular importance to select only NPCs that gave rise to neurons competent to show detectable Aβ and phosphorylated tau puncta in the neurites within the average duration of the experiments, in basal conditions. In these neurons, RNAi interference was applied for a week (from DIV4 to DIV11) to trigger a prolonged knockdown of the gene candidate. The window was tailored to the features of both lines, in a stage where mature markers are expressed but preceding the peak of production of Aβ or phosphorylated tau. The effects of gene KD were analysed using a panel of readouts: (i) levels of secreted Aβ oligomers in the supernatant by a 6E10-6E10 homotypic ELISA, (ii) intracellular/membrane associated Aβ and phosphorylated tau puncta or clusters, using monoclonal W02 and AT8 and polyclonal pSer396 antibodies and high-content confocal microscopy (HCCM), (iii) thickness, area and distribution of axons and dendrites, using MAP2 and Tuj1 antibodies and HCCM, (iv) levels of synapsin-1 puncta, also assessed by HCCM, and (v) electrophysiological activity, by longitudinally following the electrophysiological activity of neurons plated onto MEA plates. Lastly, a modification of the culture conditions of neurons subjected to candidate KD was tested. After DIV14, half-medium changes were replaced by top-up with fresh medium and concentrated factors, also containing human astrocyte conditioned medium. Levels of APP expression, Aβ oligomers, intracellular perinuclear and neurite Aβ puncta, and phosphorylated tau clusters were explored in these conditions using western blot, 6E10-6E10 homotypic ELISAs, HCCM and STED microscopy (see below).

The number of biological and technical replicates of each independent experiment is reported in the respective figures. The screening in SH-SY5Y cells was run once per gene candidate, including at least 3-5 technical replicates per condition. Experiments ran in hiPSC-derived neurons were repeated 2 to 3 times, unless otherwise stated, with technical replicates ranging 3 to 16, attending to the assay. We note that many of our readouts rely on immunocytochemistry and imaging. Apart from STED microscopy, we performed these assays in 96-well plate formats and applied high-content confocal microscopy for image acquisition. At least 12 up to 25 fields were acquired per well or technical replicate. Automated image acquisition removed user bias and allowed a better representation of the whole sample.

### Statistical analysis

Statistical analysis tailored to each experiment was performed using GraphPad Prism version 10. To compare results between ICC quantifications or ELISA readouts among the tested genes in a particular line, we performed a one-way analysis of variance (ANOVA), with Bonferroni’s multiple comparison test. When comparing the ±KD effect across different time points or medium change regimes, we performed a two-way ANOVA with Tukey’s correction for multiple comparisons.

## Acknowledgments

We thank Dr Jean-Baptiste Sibarita, IINS, Bordeaux for sharing PALMTracer analysis tool. STED imaging was performed using the STED microscope funded by BBSRC grant BB/R000395/1 at the Department of Genetics, School of Biological Sciences, University of Cambridge. We thank Dr Martin O. Lenz and Dr Antonina Kruppa for support and help. Deconvolution was performed using the Light Microscopy Core Facility at Cancer Research UK Cambridge Institute. We thank Andreas Bruckbauer and Fadwa Joud for their support at the Imaging Core Facility at the Cancer Research UK Cambridge Institute.

