## Supplementary Information for "A human neuronal model of sporadic Alzheimer’s disease induced by *FBXO2* downregulation shows Aβ aggregation, tau hyperphosphorylation and functional network impairment"

### Materials and Methods

**Sourcing and handling siRNAs.** Pre-designed Silencer Select siRNAs against each target were procured from Ambion (Thermofisher). A 10 µM siRNA stock was prepared by mixing at least two siRNAs duplex sequences targeting two different regions of each mRNA target. Silencer™ Select Negative Control No. 1 siRNA duplexes (#4390843, Ambion, Thermo Fisher) were used as a negative control. siRNA duplexes were kept at -20 °C and thawed in ice before use.

**Screening of gene candidates in SH-SY5Y cells. *Culture and maintenance.*** Human neuroblastoma SH-SY5Y cells were authenticated and subjected to mycoplasma testing before initiating the experiments. Cells were cultured in maintenance medium (MM, DMEM/F-12 Glutamax™, #10565018, Gibco), supplemented with 10% v/v of heat-inactivated fetal bovine serum (FBS, #10082147, Gibco) and kept at 37 °C, 5% $CO_2$ and 95% humidity levels, unless otherwise indicated. Passage number was kept below 20 for all performed assays. For gene candidate screening, cells were detached using Trypsin-EDTA (0.25%) solution (#25200056, Gibco) and replated in PerkinElmer Cell Carrier Ultra 96 well plates at a density ranging from 12.5k to 15k cells/well in MM, with a final volume of 100 µL/well. Cells were incubated for at least 24 h to allow adequate attachment prior RNA interference (RNAi). ***RNA interference.*** 1 h before RNAi, the cell medium was replaced by the transfection medium (TF, DMEM/F-12 Glutamax™ supplemented with 1% v/v FBS), with a final volume of 90 µL per well. siRNA duplexes were prepared a maximum of 15 min before their addition to the supernatant. In brief, siRNAs and lipofectamine RNAiMax (#13778150, Invitrogen) were diluted in Opti-MEM reduced serum medium (#31985062, Gibco) following the manufacturer´s specifications. siRNA and lipofectamine solutions were mixed at a 1:1 ratio and incubated for 5 min at RT. 10 µL of the lipo-complexes were added per well, reaching a final concentration of 10 nM per well. ***Exogenous treatment with Aβ monomers.*** 16 h after transfection, the medium was discarded and replaced by 75 µL of fresh TF. Cells were incubated for 6-8 h at 37 °C and treated with 25 µL of freshly purified monomeric Aβ(M1-42) at a final concentration of 2-4 µM. Cells were then incubated for 1 to 3 days at 37 °C, to provide enough time for the Aβ monomers to get internalised and aggregates to build-up. Every 24 h, the cells were fixed with 4% w/v PFA (#28906, Thermo Scientific) in D-PBS (+/+) (#14040141, Gibco) and stained with the W0-2 Aβ-specific antibody (#MABN10, Sigma Aldrich, 1:500) to study the presence, distribution and morphology of Aβ aggregates. Since the time scale over which Aβ nucleates in cellular membranes and becomes internalised in cultured cells is variable between different peptide batch preparations, we always collected three time points (24, 48 and 72 h) after each Aβ treatment. We then analysed and reported data from the earliest time point in which Aβ aggregation is detectable in our negative control, ScrRNA-treated cells. This result indicated that the Aβ treatment was successful and basal aggregation could be observed, even without any gene expression perturbation.

**hiPSCs culture and maintenance.** Two human induced-pluripotent stem cell (hiPSCs) lines were used in the present study: (a) the BIONi010-C-38 line, referred to as $APP^{Swe}$ in this work, obtained from Bioneer, and (b) the Kolf2-C1 line, referred to as $APP^{WT}$, obtained from the Wellcome Sanger Institute. Cells were authenticated and tested for mycoplasma contamination before any experimental application. hiPSCs were maintained as previously reported (*1*). In brief, hiPSC colonies were cultured in GelTrex™-coated flasks (#A141320, Gibco). Cells were grown in mTeSR Plus (#100-0276, StemCell Technologies) and split using ReLeSR™ (#100-0484, StemCell Technologies) when colony densities went above 70%. Medium change was changed every other day, unless otherwise indicated.

**hiPSCs differentiation to cortical NPCs.** hiPSCs were differentiated into cortical progenitor cells by adapting a previously reported protocol (*2*), as already described in detail (*1*). In brief, the protocol involved breaking hiPSC colonies into single cells, plating them to full confluency in GelTrex™-coated flasks and inducing neural differentiation using neural maintenance medium (NMM) supplemented with different small molecules and morphogens. The neural maintenance medium (NMM) was prepared by mixing 1:1 a N-2 and B-27-containing media. The N-2 medium was prepared with DMEM/F-12 Glutamax™, 1X N-2 (#17502048, Gibco), 5 µg/mL of human insulin (#I9278, Sigma), 100 µM of non-essential amino acids (#11140050, Gibco) and 100 µM 2-mercaptoethanol. THe B-27 medium was prepared with Neurobasal (#21103049, Gibco) and 1X B-27 (#17504044, Gibco). To form a neuroepithelial layer, the culture was treated with 10 µM SB431542 (#S4317, Sigma Aldrich) and 100 nM LDN-193189 (#SML0559, Sigma Aldrich) in NMM for 12 days, followed by manual procedure consisting of breaking the neuroepithelium into 100-200 µM clumps, which are subsequently replated in fresh GelTrex™-coated flasks for rosette induction. Rosette development is then triggered using 1 µg/mL of CHIR99021 (#SML1046, Sigma Aldrich) and 20 ng/mL of FGF2 (#3718-FB, RnD Systems) until their cryopreservation on days 20-22.

**Differentiation of cortical NPCs to glutamatergic neurons.** Cortical NPCs were thawed on GelTrex™-coated 6-well plates in NMM at 100 % confluency. The medium was refreshed for 3 consecutive days. At day 4, neural progenitors were collected with the use of Accutase (#A6964, Sigma Aldrich) and dislodged into single cells using NMM enriched with ROCK inhibitor (#SCM075, Sigma Aldrich) and 100 U/mL of penicillin and streptomycin (#15140122, Gibco). Cell suspension was then distributed into PerkinElmer CellCarrier Ultra 96 well plates pre-coated with 0.002% poly-L-ornithine (#P4957, Sigma Aldrich) and 10 µg/mL of laminin (#L2020, Sigma Aldrich), at a density of 90-100k cells/well. The culture medium was refreshed with NMM supplemented with 10 ng/mL of rhBDNF (#248-BDB, R&D Systems), 10 ng/mL of rhGDNF (#212-GD, R&D Systems), 200 µM of L-ascorbic acid (#A4544, Sigma), 500 µM of cAMP (#D0627, Sigma), 1X of CultureOne (#A3320201, Gibco) and 100 U/mL of penicillin-streptomycin (Pen/Strep) on days 1, 4 and 7 post-seeding. Cells were also exposed to 10 µM DAPT (#2634/10, Tocris) on days 1 and 4. Starting on day 11, the medium was changed every two to three days using BrainPhys™ Neuronal Medium (#05790, StemCell Technologies) supplemented with 1X N-2, 1X SM1 (#05711, StemCell Technologies), 10 ng/mL of BDNF (#248-BDB, R&D Systems), 10 ng/mL of GDNF (# 212-GD, R&D Systems), 200 µM of L-ascorbic acid (#A4544, Sigma), 500 µM dibutyryl cyclic-AMP (#D0627, Sigma Aldrich), 1X CultureOne™ (#A3320201, Gibco), and 1X Pen/Strep.

**Quality controls of NPCs and derived cortical glutamatergic neurons.** First, to assess NPCs phenotypic identity, purity and potency, cryopreserved progenitors were thawed and plated into GelTrex™-coated PerkinElmer Cell Carrier Ultra 96-well plates at a density of 100k/well using 1X N2B27 supplemented with Pen/Strep and Rocki. NPCs are incubated for 24 h at 37 °C prior fixation with 4% PFA solution, prepared in D-PBS (+/+) (DIV0). To assess the phenotype and purify of NPC-derived neurons, NPCs were thawed and replated into PLO and L2020-coated PerkinElmer Cell Carrier Ultra 96 well plates.

We then conducted thorough quality checks using immunocytochemistry to measure the levels of the following markers: EMX1, DLX5, NXK2.1, PAX6, Ki67, OTX1/2, SATB2, nestin and TBR1. For the NPCs to be considered suitable for use, we required that 60-80% of the cells tested positive for EMX1, DLX5, PAX6, OTX1/2, SATB2, nestin and TBR1 at early stages of the terminal differentiation protocol (from DIV0 to DIV4). These markers are indicative of progenitor cells destined to become dorso-anterior, cortical, excitatory, glutamatergic neurons (*3-7*). Additionally, the good quality NPCs were defined low levels of NXK2.1 –a transcription factor enriched on progenitors of ventral forebrain interneurons (*3*) and Ki67 – an indicator of active cell division (*8*).

**Gene candidate screening in glutamatergic neurons.** At DIV4, DIV7 and DIV11, cortical neurons were treated with 5 to 10 nM siRNA duplexes against each gene of interest, attending to the cell density in the well. siRNA and lipofectamine RNAiMax complexes were prepared as described in sections above. For the treatment, 10 µL of medium per well was removed and replaced by 5 or 10 µL of the siRNA lipocomplex solution. No medium change was performed following transfections.

**Cellular lysis for RNA extraction.** The cell medium was carefully removed from the corresponding wells. Afterwards, 50-100 µL of lysis buffer (#12183025, Invitrogen) freshly enriched with 1% β-mercaptoethanol (#M6250, Sigma Aldrich) were added per well, followed by 5-10 min incubation at room temperature. Lysates from at least 4 wells were combined into a single RNAse free Eppendorf tube, which were subsequently flash-frozen and stored at -80 °C for further RNA purification.

**RNA purification.** Cell lysates were thawed at room temperature until a clear solution was observed. 400 µL of 70% EtOH were added to each 400 µL of cellular lysates. Tubes were gently inverted 10 to 15 times, until obtaining a clear solution. A maximum of 700 µL of the previous solution was transferred into labelled RNA spin column tubes (#12183025, Invitrogen). Purification was performed as per manufacturer's protocol.

**Messenger RNA analysis by quantitative PCR (qRT-PCR).** 250 ng to 1 µg of RNA were used to synthesize cDNA in a 20 µl volume with the High-Capacity RNA-to-cDNA kit (#4388950, Applied Biosystems). The quantitative real-time polymerase chain reaction (qRT-PCR) was carried out in a volume of 20 µL using the TaqMan™ Gene Expression Master Mix (#4369016, Applied Biosystems) and 1X of each specific TaqMan probe (Thermo Fisher). An initial 10 min denature step of 2 min at 50 °C and 10 min at 95 °C were followed by 40 cycles of denaturation (15 sec at 95 °C), annealing (30 sec at 60 °C) and elongation (30 sec at 60 °C). Both retro-transcription and qPCR reactions were run in a QuantStudio3™ Real-Time PCR System (Applied Biosystems, UK). Relative expression levels of our genes of interest were evaluated with the ΔΔCt method, using ACTB or GAPDH as housekeeping genes, whose expression is determined to be stable between experimental conditions.

**6E10-6E10 Capture-Detector ELISA for quantification of secreted Aβ oligomers.** Streptavidin-coated 96 well plates (#436014, Thermo Scientific) were washed with 300 µL per well of 0.05% Tween-20 in PBS. Afterwards, each well was coated with 70 µL of a 1:500 dilution of a biotinylated 6E10 antibody (#803008, BioLegend) in 0.05% Tween-20. The capture antibody was incubated for 1 h at RT under shaking conditions (250 rpm). Cell supernatants were thawed at RT and centrifuged at 4000 rpm for 15 min at 4 °C to remove insoluble aggregates and cellular debris. A protease inhibitor solution was added at 1X final concentration to each sample (COEDTAF-RO, Roche). The capture antibody was washed twice with 0.05% Tween-20 in PBS before sample addition. Samples were incubated for 2 to 3 h at 4 °C under shaking conditions (250 rpm). The plates were then washed three times with TBS prior addition of HRP-6E10 detector antibody (#803012, BioLegend) diluted in a 5% BSA solution in TBS at a concentration of 0.1 µg/mL. The detection antibody was incubated 1 h at RT under agitation (250 rpm). Plates were washed three times in TBS. Afterwards, 100 µL of TMB ELISA (#34028, Thermo Scientific) substrate was added per well. The reaction was left to develop for 30 min in the dark after being stopped with the addition of 50 µL per well of 2M HCl. Absorbance values were measured at a wavelength of 450 nm using a plate reader (BMG Labtech, Aylesbury, UK).

**Differentiation of NPCs to neurons in MEA plates and recording of electrophysiological activity.** ***Plating and differentiation protocol.*** Cortical NPCs were thawed on GelTrex™-coated 6-well plates in NMM at 100% confluency. The medium was refreshed for three consecutive days before replating on MEA plates. 24 h prior replating, CytoView MEA 96 well plates (Axion Biosystems) were coated with 80 µL per well of 0.07% PEI solution (#P3143, Sigma Aldrich) prepared in 1X Borate Buffer (#28341, ThermoScientific). MEA plates were incubated at 37 °C for 1 h. Plates were then washed twice with > 260 µL per well of sterile D-PBS and once with the same volume of sterile water. Special care was taken during the manipulation of the MEA plate, avoiding scratching the surface of the electrode with the pipette tips. Water was then aspirated, and plates were left to air-dry overnight inside a sterile biological safety cabinet. NPCs were collected and counted as reported in the section "differentiation of cortical NPCs to glutamatergic neurons". 14.4 million cells were centrifuged and resuspended in 1.2 mL of dotting medium (100 µg/mL of L2020 in 1X N2B27 and 10 µM Rock inhibitor) for a final concentration of 12 million cells/mL, avoiding the generation of bubbles. 10 µL droplets from the cell suspension were dispensed precisely on the center of each well, covering all the electrodes. Sterile warm water was added in the surrounding plate reservoirs to prevent evaporation and the plate was incubated at 37 °C for 1 h. 200 µL of 1X N2B27 supplemented with Rock inhibitor was added carefully per well, avoiding dislodging adhered NPCs. NPCs were differentiated to mature cortical glutamatergic neurons following the protocol described in the section "differentiation of cortical NPCs to glutamatergic neurons", by performing half-medium changes three times per week. ***Recording.*** We recorded spontaneous electrical network activity at desired time points was recorded by performing a 20-min equilibration period, followed by one measurement of 10 min. All recordings were conducted using the Axion Integrated Study (AxIS) software, under a controlled environment of 37 °C and 5% $CO_2$. The electrical activity was measured with a gain of 1000x and a sampling frequency of 12.5 kHz. Before spike detection, a Butterworth band-pass filter ranging from 100 to 3000 Hz was applied. Spike detection was performed using the AxIS adaptive spike detector, with a threshold set at 6 times the root mean square (RMS) noise on each electrode. An electrode was considered active if it exhibited a spike rate of at least 5 spikes/min. These recording and analysis parameters were employed to evaluate and characterize the spontaneous electrical network activity in the experimental setup. Each recording was re-recorded using the AxIS software, generating .spk files. Network bursts were identified using the Axion Neural Metric Tool. These detection methods yielded 57 parameters related with individual neuronal activity, network burst and network synchrony.

**Immunocytochemistry**
Cells were fixed with a solution of 4% PFA diluted in D-PBS (+/+), at room temperature for 10 min. Fixed cells were then washed three times with D-PBS (+/+), followed by permeabilization with 0.1% v/v of Triton X-100 in D-PBS (+/+) (#85111, Thermo Scientific) for 30 min at RT. Afterwards, cells were treated with immunofluorescence blocking buffer (#12411, Cell Signalling Technology) for 1h at RT. Blocking solution was removed and cells were incubated with the appropriate primary antibody solution overnight at 4 °C. After four washes with D-PBS (+/+), cells were then incubated with AlexaFluor-conjugated secondary antibodies for 1h at room temperature in the dark. Both primary and secondary antibodies were diluted in immunofluorescence blocking solution. Cells were wash other four times with D-PBS (+/+). Nuclei was stained using Hoechst 33342 (#H3570, Thermo Fisher) at 1:1000 dilution, also in D-PBS (+/+). The list of primary and secondary antibodies used in this study is detailed in **Table S2**.

**Treatment with astrocyte conditioned medium**
$APP^{WT}$ and $APP^{Swe}$ cortical NPCs were thawed and replated to 96 well plates as described in previous sections. RNA interference was performed from DIV4 to DIV11, as reported above. Until DIV14, cells were cultured in 100 µl of medium. At DIV14, 50 µl were removed per well and replaced with 20 µL of either BrainPhys (NM) or a 1:1 mixture of BrainPhys and Astrocyte Conditioned Medium (#1811, ScienCell) (ACM), in both cases supplemented with 35 ng/mL BDNF, 35 ng/mL GDNF, 700 µM

ascorbic acid, 1.75 mM of cAMP, 3.5X CultureOne™ and 3.5X Pen/Strep. From DIV14 onwards, a top-up medium change regime was applied, meaning that no medium was removed from the wells until the completion of the experiment. 20 µl per well of NM or ACM supplemented with BDNF, GDNF, AA, cAMP, CultureOne™, Pen/Strep were performed every two to three days, as detailed in **Figure S6**.

**Western blot**

Protein extraction was performed by lysing cells directly in the wells using N-PER neuronal protein extraction reagent (#87792, Thermo Scientific) supplemented with phosphatase and protease inhibitors (#78440, Thermo Scientific), as per manufacturers protocol. Lysate was centrifuged at 10,000 x g for 15 minutes at 4°C to pellet cellular debris, and the supernatant containing soluble proteins was collected and stored at -80°C until further analysis. Protein concentration was determined using the Pierce BCA Protein Assay Kit (# 23225, Thermo Fisher Scientific), with bovine serum albumin (BSA) used to generate a standard curve. Samples and standards were prepared at least in duplicate, with 25 µL of each sample or standard added to a 96-well plate. Subsequently, 200 µL of working reagent (prepared according to the manufacturer's instructions) was added to each well, and the plate was incubated at 37°C for 30 minutes. Absorbance was measured at 562 nm using a microplate reader.

For SDS-PAGE, protein lysates (12.5 to 30 µg per lane) were mixed with NuPAGE™ LDS sample buffer (#NP0007, Invitrogen) supplemented with NuPAGE™ sample reducing agent (#NP0004, Invitrogen) and boiled at 95°C for 10 minutes. Samples were then loaded onto a 4–12% NuPAGE™ Bis-Tris Mini Protein Gel (#NP0321PK2, Thermo Scientific). Running parameters were set for 35 min at a constant 200 V. Following electrophoresis, proteins were transferred to nitrocellulose membranes using an iBlot2 Dry Blotting System (Thermo Fisher Scientific) with a transfer program set to 7 minutes.

After transfer, the membrane was blocked in 5% non-fat dry milk in Tris-buffered saline with 0.05% Tween-20 (TBS-T) for 1 hour at room temperature. The membrane was then incubated with the corresponding primary antibody diluted in 5% non-fat dry milk in TBS-T overnight at 4°C. Following primary antibody incubation, the membrane was washed three times for 10 minutes each with TBS-T and subsequently incubated with the HRP-conjugated secondary antibody diluted in TBS-T with 5% non-fat dry milk for 1 hour at room temperature. After secondary antibody incubation, the membrane was washed three times for 10 minutes each with TBS-T. Protein bands were visualized using an enhanced chemiluminescence (ECL) substrate (Thermo Fisher Scientific) and imaged using a ChemiDoc Imaging System (Bio-Rad).

**Image acquisition and analysis**

Unless otherwise stated, image acquisition was performed using Opera Phoenix High-Content Confocal microscope using a water 20X or 40X objective. At least 12 images per technical replicate for each biological replicate were taken for quantification. Image analysis and quantifications were performed with the Harmony High-Content Imaging and Analysis Software (Perkin Elmer).

**Stimulated emission depletion microscopy (STED) and cluster analysis**

We used a commercial STED inverted microscope (Abberior STED Expert Line Super Resolution Microscope, Abberior Instruments GmbH, Göttingen, Germany). This laser scanning microsocope provides multiple super-resolution and confocal channels. It is based on a fully automated Olympus IX83 microscope platform with a 100x (1.4 NA) oil immersion objective (UPLSAPO 100XO) and a high-precision Ultrasonic Stage (IX3-SSU). STED and confocal image acquisition was performed with pulsed 488 nm, 561 nm and 640 nm laser excitation and depletion at 775 nm. Images were taken for a region of interest (750 pixels x 750 pixels) with a sampling of 20 nm. The pinhole was set to 1.0 AU.

For dual-colour STED imaging, the same depletion laser was used in lateral depletion mode while the excitation wavelength was adapted to the fluorophore. The Alexa Flour 488 images were recorded only in the confocal mode. The image acquisition was controlled by Imspector software.

Deconvolution was performed using Huygens Professional software (Scientific Volume Imaging, Hilversum, Netherlands) in STED mode, employing the Classic Maximum Likelihood Estimation (CMLE) algorithm. Clusters of molecular aggregation were identified from super-resolution STED images by a custom algorithm written as a plug-in supported by MetaMorph (Molecular Devices). Clusters were detected from super-resolution images using PALMTracer plugin (*9-11*) (Kechkar 2013, Izeddin 2012, Butler 2022) and analyzed as described previously (*12-14*) (Kedia 2021, Kedia 2020, Kedia 2020).

**Compilation of a list of genes linked with early transcriptomic perturbations in AD**

To study early transcriptomic perturbations in sAD, we used snRNA-seq data from the prefrontal cortex of 48 participants in the ROSMAP longitudinal cohort studies of ageing and dementia (*15*). We performed K-means clustering based on the clinico-pathological readouts available to determine a classification in 24 healthy individuals, 15 early-AD and 9 sAD patients (**Figure S7A**). Broadly consistent with the NIA-AA guidelines (*16*), the distinction between early AD (fAD) and late AD (sAD) stages was driven by higher neurofibrillary tangles (NFTs) burden and density and lower cognitive function, whilst a comparably high amyloid burden was observed overall among the stages (**Figure S7A**).

To study the cell-type specific disruption to the biochemical pathways in AD, we initially separated main cell types as previously described (*15*), and a set of marker genes *(17).* Using a uniform manifold approximation and projection (UMAP) approach, we identified eight cell types: excitatory neurons (Ex), inhibitory neurons (In), astrocytes (Ast), microglia (Mic), oligodendrocytes (Oli), oligodendrocyte progenitor cells (Opc), pericytes (Per), and endothelial cells (End) (**Figure S7B,C**). We did not identify any relevant cell classification based on the pathological staging of the disease, owing both to the subtlety of the molecular perturbations occurring in AD, which cannot be singled out from a low-dimensional representation of the transcriptome, and to the presence of subpopulations of cells within each different cell type (*17, 18*)*,* that can respond differently to the disease insults to the brain (*19, 20*). We further separated each cluster of cells into subpopulations, using the Louvain graph-clustering method (*21, 22*), detecting a cell-type dependent heterogeneous number of sub-clusters, proportional to the number of nuclei populating the main clusters (**Figure S8**).

We then applied Pathifier (*23*), a method of analysing the overall expression of biochemical pathways and assigns a dysregulation score that quantifies the extent of their perturbation in disease with respect to physiological conditions. Originally developed to improve the stratification of disease samples in cancer by studying dysregulation of known biochemical pathways perturbed in disease, we used this approach to identify the top perturbed pathways in AD, building on the stratification of the samples described above (**Figure S9**). We applied the method on all the 324 pathways in the KEGG annotation (*24*) across the cell subpopulations described above. An extremely heterogeneous scenario of perturbations emerged from this analysis, with neuronal cell types undergoing widespread dysregulation in multiple pathways. Excitatory neurons amongst the neuronal types, and oligodendrocytes and astrocytes within the glia cells were the most vulnerable to pathway-based perturbations (**Figure S10A**). Despite this widespread vulnerability, perturbations on neurons seemed to be more pronounced at the early stages of AD, while for glia cells the dysregulation peaked later during its progression (**Figure S10B**).

Considering the high heterogeneity of the perturbations, we studied a subset of KEGG pathways associated with AD, called network-based transcriptome-wide association study (nTWAS) pathways, thought the Pathifier analysis (**Figure S7D,E**). We measured in how many of the cell subpopulations the nTWAS pathways were found to be significantly perturbed, and observed that these pathways were consistently more impaired than the rest, particularly in the neuronal and astrocytes populations. They were found to be highly perturbed both at early and late stages of AD (**Figure S7D,E**). Moreover, the relevance of nTWAS pathways in neurodegeneration was supported by studying the correlation between their transcriptomic dysregulation and the measured effect on AD clinical manifestations during disease progression. The nTWAS set was highly correlated with all the major clinical traits associated with AD, and strongly anti-correlated with cognitive functions in turn, without showing any significant interdependence with ageing (**Figure S11**). These results denote a strong specificity of these pathways to neurodegenerative processes apart from that pertaining to general ageing, as they could disentangle sAD progression from its major risk factor and confounder (**Figure S11**).

We then investigated all the pathways able to discriminate between different pathological states into more details. The KEGG pathways were initially clustered based on their similarity, intended as the genes overlap between them to determine whether a fingerprint of the dysregulation could be detected in specific groups. We initially identified 27 clusters from the 324 KEGG pathways (**Figure S12**), 3 of which were significantly enriched in nTWAS pathways (**Figures S13 and S14**). To decipher the heterogeneity at the subpopulations level, we then performed a consensus analysis and considered a perturbation of a pathway to be statistically significant only if the combined p-value across sub-clusters within a cell type also resulted to be significant. From the widespread dysregulation, our results showed that the nTWAS pathways, and their clusters of reference, were especially overrepresented in neuronal cells (**Figure S14**). Several nTWAS pathways were also recurring in glia cells, with the cluster enriched in neurodegenerative disorders (cluster 2 in **Figure S13**) being the most recurrent across all cell types (**Figure S14A**), supporting the robustness of the Pathifier approach towards our application in AD. In general, we found that nTWAS-enriched clusters 16 and 18 were the most significantly dysregulated in neuronal cells.

The evidence of the centrality of nTWAS pathways in neurodegeneration led us to investigate how the perturbations to the pathways involved changed across disease stages. We observed a stage-dependent shift in the impact of AD on the signalling pathways and the ones belonging to the neurodegeneration cluster. In fact, signalling and synaptic transmission pathways, majorly impaired in early AD, collectively dropped in terms of relevance in later AD stages, while inflammation and neurodegenerative processes cooperatively emerged (**Figure S7E**). This shift in importance of the signalling pathways may be interpreted in terms of the formerly reported impairment in the excitation-inhibition balance causing neuronal hyperactivity at early stages in AD (*25*).

As signalling processes appeared to be the most relevant at early stages, to single out the major contributors to the nTWAS pathways perturbations, we ranked the genes involved according to their overall importance in determining the perturbation trajectories. The sub-clusters of the neuronal cells and astroglia in which the nTWAS processes were found to be significantly perturbed were then selected, and their genes ranked to perform a consensus analysis. For each pathway the rankings were combined across the relevant subpopulations by summing them to define an overall classification of the most important genes. This allowed to determine which genes would consistently rank higher across subclusters and contribute more to define the dysregulation trajectory. Among the top contributors to pathways perturbations which were also differentially expressed in early AD in neuronal cells or astroglia, we found several subunits (i.e. ATP6V0B, ATP6V0E2, ATP6V1B2, ATP6V1E1, ATP6V1F, and ATP6V1H, with the last one being only highly perturbed but not differentially expressed) of the two complexes composing the vacuolar-type ATPase (v-ATPase), a multimeric protein complex which acts as an ATP-dependent proton pump and controls the acidic pH of the lysosome. Lysosomal dysfunction and more specifically dysregulation of the lysosomal acidification have been linked both to tau-induced toxicity, and to failures in degradation of Aβ fibrils in animal models of tauopathies and AD, respectively (*26-28*). Furthermore, the expression levels of RAB3A, a small GTP-binding protein key in regulating exocytosis and secretion, as well as in the assembly of vesicles deputed to the anterograde transport of APP (*29*) and for regulating lysosome exocytosis (*30*) resulted to be significantly lowered in cortical brain regions of AD patients (*31, 32*). Another 4 genes identified, namely CLTA, CLTB, and CLTC, encoding two light chains and one heavy chain of clathrin, respectively, and FLOT1, an integral membrane protein, are all positive regulators of endocytosis; their measured centrality in pathway-based perturbations, together with their downregulation in early AD (*15*) denote how reduced vesicles recycling could play a major role in the onset of AD, as well as in its early progression. In fact, in a recent work, an early intervention strategy targeting the regulation of the dysfunction of CLTC-mediated endocytosis for a synaptic receptor showed promising results (*33*),

Moreover, in agreement with our findings, RAB3A, together with proteins involved in the synaptic vesicle cycle at the stage of docking and proteins priming (e.g. SNAP25 and VAMP2, two SNARE proteins involved in vesicle docking, membrane fusion, and exocytosis) were recently identified as potential biomarkers for detection of early signs of AD. Their altered abundances derived from in-depth proteomics of the prefrontal cortex of AD patients (*34, 35*) highlight their potential role in diagnostics and early therapeutic intervention (*36*). Remarkably, the strong correlation existing between intracellular Aβ and the progression of AD symptoms in the early stages of AD indicates that the presynaptic dysfunction induced by its intracellular fraction might be one of the pathophysiological origins of early AD (*37*). This is in line with the centrality of the synaptic vesicle cycle in the aetiology of AD.

Taken together, these observations shed a light on a possible mechanistic interpretation of how a progressive disruption of the synaptic vesicle cycle, at the level of the endosomal-lysosomal machinery and exocytotic events, could represent an additional hallmark of the early stages of Alzheimer's disease (*38*). This list of gene dysregulated in early AD is reported in **Figure 1A**.

**Table S1. List of shortlisted genes from the SH-SY5Y screening assay**.

| Shortlisted genes from SH-SY5Y screening assay |
|---|
| *ATP6V0E2* |
| *ATP6V1B2* |
| *ATP6V1E1* |
| *ATP6V1G1* |
| *CHMP5* |
| *CTLA* |
| *CTLB* |
| *CTLC* |
| *RAB3A* |
| *RAB11A* |
| *SLC17A7* |
| *SORCS3* |
| *USF3* |
| *VPS35* |
| *VPS36* |
| *FBXO2* |

### Table S2. List of antibodies used in this work

| **PRIMARY ANTIBODIES** | | | |
|---|---|---|---|
| TARGET | HOST, ISOTYPE, CLONE | CATALOG NUMBER | WORKING DILUTION |
| Anti-APP/Aβ | Mouse, IgG2aκ, W02 | MABN10, Sigma Aldirch | 1:500 |
| Aβ40 | Mouse IgG1, κ | 805409, BioLegend | 1:50 |
| Aβ42 | Mouse IgG1, κ | 805509, BioLegend | 1:50 |
| Tau | Mouse, IgG1, T46 | 13-6400, ThermoFisher | 1:250 (ICC) |
| Tau | Chicken, IgY | NBP2-25163, BioTechne | 1:1000 (STED) |
| Phospho-Tau (Ser202, Thr205) | Mouse,IgG1κ, AT8 | MN1020, ThermoFisher | 1:1000 |
| Phospho-Tau (Ser396) | Rabbit, IgG | 44-752G, ThermoFisher | 1:1000 |
| Phospho-Tau (Thr212) | Rabbit, IgG | 44-740G, ThermoFisher | 1:1000 |
| EMX1 | Rabbit, IgG | PA5-35373, ThermoFisher | 1:50 |
| DLX5 | Rabbit, IgG, EPR4488 | ab109737, Abcam | 1:1000 |
| NKX2.1/TTF1 | Rabbit, IgG, EP1584Y | ab76013, Abcam | 1:500 |
| Ki67 | Mouse, IgG1 κ | 550609, BD Biosciences | 1:600 |
| Nestin | Mouse, IgG1, 196908 | ab6320, Abcam | 1:1000 |
| PAX6 | Rabbit, IgG | 42-6600, Invitrogen | 1:500 |
| OTX1/2 | Rabbit, IgG | AB9566-I, Sigma Aldrich | 1:250 |
| SATB2 | Rabbit, IgG | PA5-19630, Invitrogen | 1:300 |
| TUJ1 | Mouse, IgG2aκ | 801201, Biolegend | 1:1000 |
| TUJ1 | Rabbit, IgG | ab52623, Abcam | 1:1000 |
| TUJ1 | Chicken, IgY | ab41489, Abcam | 1:1000 |
| MAP2 | Chicken, IgY | ab5392, Abcam | 1:3000 |
| Synapsin 1 | Mouse, IgG1, 7H10G6 | MA5-31919, Invitrogen | 1:1000 |
| PSD-95 | Rabbit, IgG | ab18258, Abcam | 1:1000 |
| VGLUT1/2 | Rabbit, IgG | 135 503, Synaptic Systems | 1:1000 |
| VGAT | Mouse, IgG3 (κ light chain), 117G4 | 131 011, Synaptic Systems | 1:1000 |
| **SECONDARY ANTIBODIES** | | | |
| ANTIBODY | CATALOG NUMBER | SUPPLIER | WORKING DILUTION |
| AlexaFluor$^{488}$ goat anti-mouse IgG (H+L) | A-11001 | Invitrogen | 1:1000 |
| AlexaFluor$^{555}$ goat anti-mouse IgG (H+L) | A-21424 | Invitrogen | 1:1000 |
| AlexaFluor$^{647}$ goat anti-chicken IgY | A-21449 | Invitrogen | 1:1000 |
| AlexaFluor$^{488}$ goat anti-rabbit IgG (H+L) | A-11008 | Invitrogen | 1:1000 |
| AlexaFluor555 goat anti-rabbit IgG (H+L) | A32732 | Invitrogen | 1:1000 |

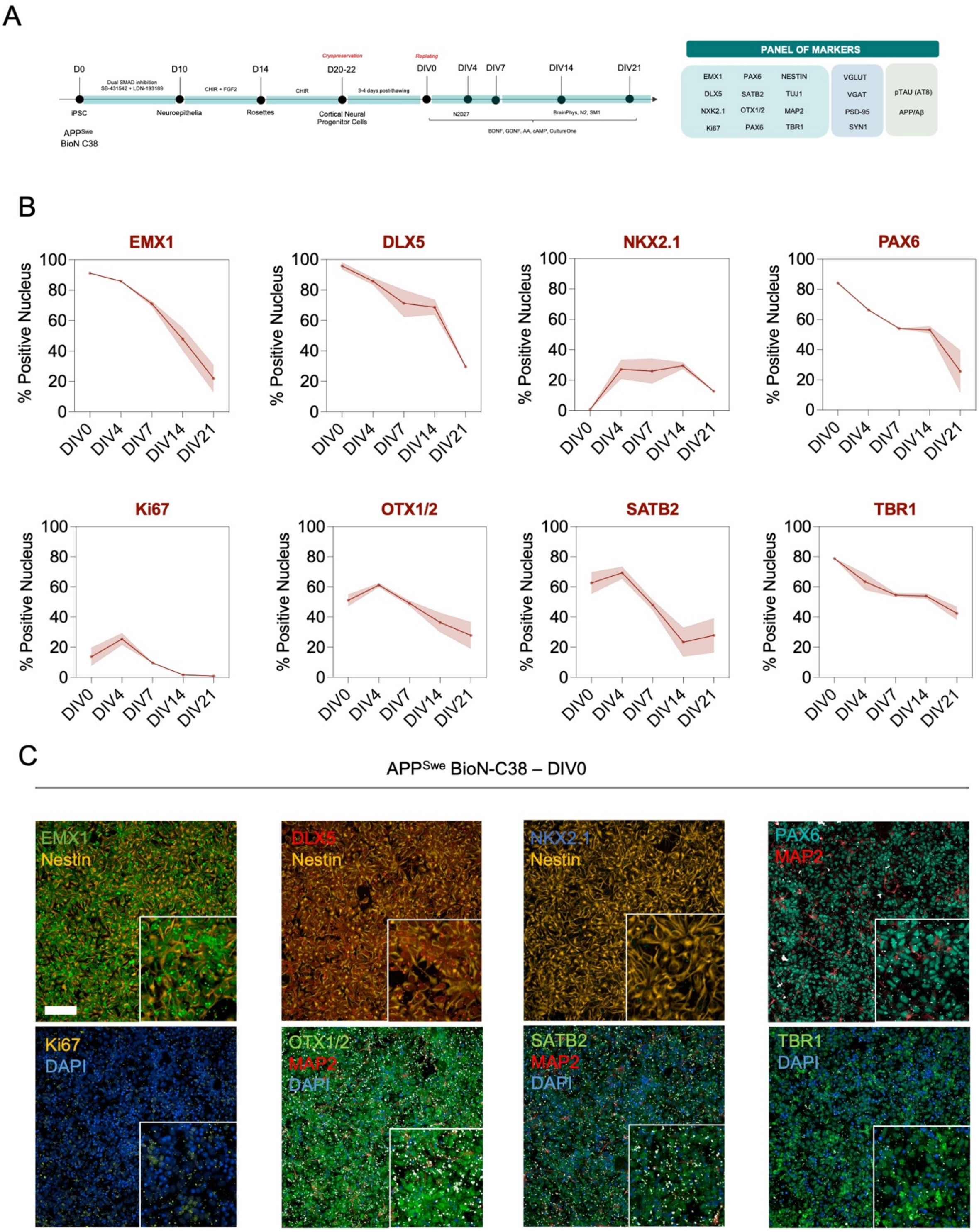


**Figure S1. Quality control of APP$^{Swe}$ cortical neural progenitor cells (NPCs) by immunocytochemistry. (A)** Illustration of the differentiation protocol used in this work, which was adapted from a previously reported one (*2*). hiPSCs are converted into neuroepithelia, followed by neural tube-like rosette formation and subsequent isolation of cortical NPCs, which then are differentiated to cortical glutamatergic neurons. The NPCs generation process takes 20-22 d. Following generation, high-purity cortical NPCs are cryopreserved in large batches to maintain population homogeneity and reduce experimental variability. **(B)** The phenotypic nature of cortical APP$^{Swe}$ NPCs is characterised by assessing their enrichment in EMX1, DLX5, PAX6, OTX1/2 and SATB2 expression, which is expected to decrease upon differentiation to mature neurons. Absence of NKX2.1 – a marker of ventral forebrain interneurons progenitors (*39*) - was also indicative of the cortical identity of our NPCs. Ki67 was used as a marker to determine the percentage of mitotic cells in culture. Data are represented as the mean and standard error of the percentage of DAPI-positive nuclei that stained positive for the expression of each transcription factor. Expression was assessed by immunocytochemistry at DIV0 (NPC stage), DIV4, 7, 14 and 21 (n = 3). (C) Representative images showcasing the expression levels of each transcription factor at DIV0. Cells were also stained for DAPI (nuclei), nestin (cytoskeleton of progenitor cells) and MAP2 (dendritic marker of neuronal maturity). Scale bar = 100 μm.

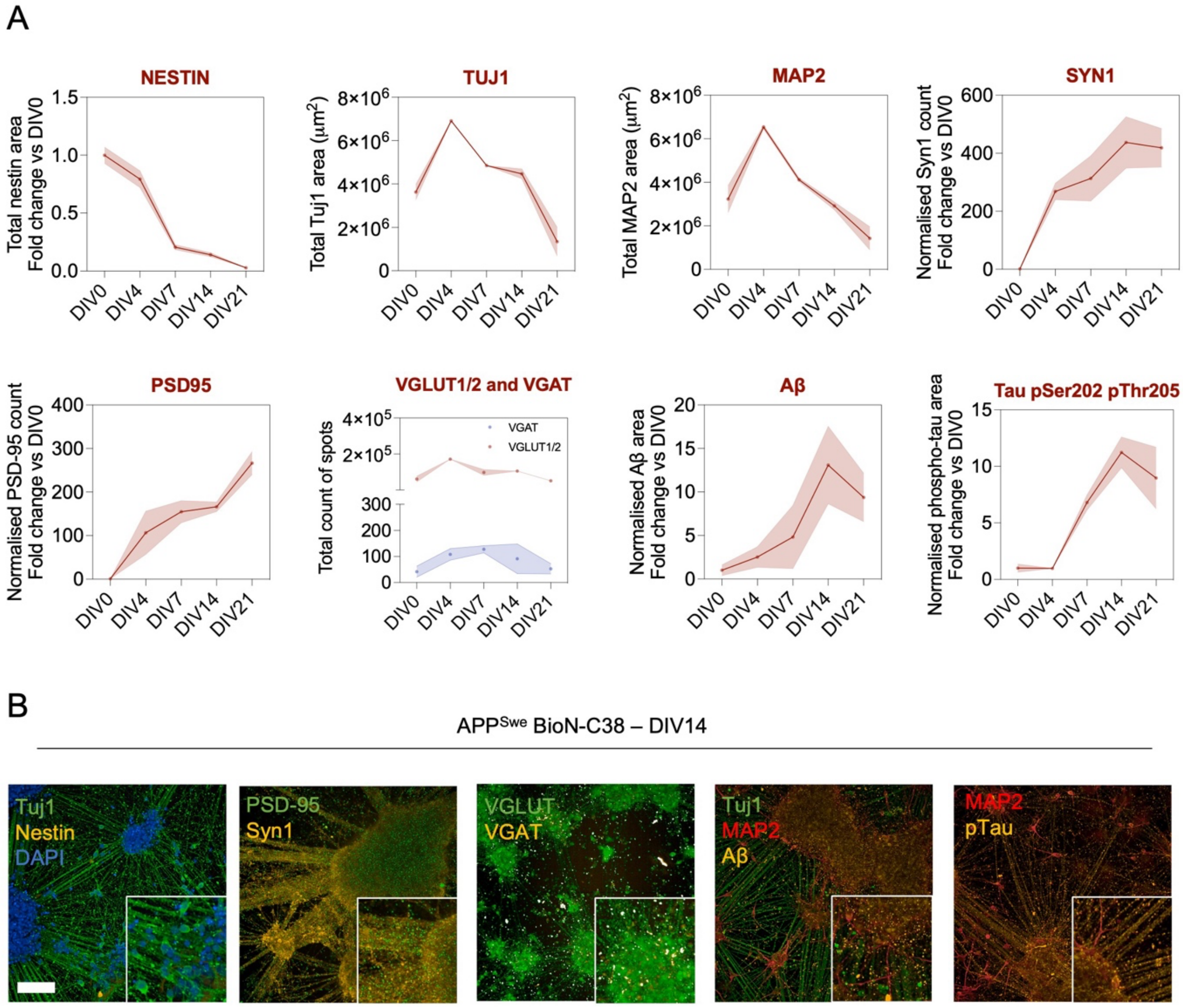


**Figure S2. Quality control of APP$^{Swe}$ cortical glutamatergic neurons by immunocytochemistry.** **(A)** The phenotypic nature of cortical glutamatergic neurons derived from the BIONi010-C-38 iPSC clone is characterised by assessing: (i) the decrease in expression levels of the progenitor-specific marker nestin upon neuronal maturation, (ii) the expression of axonal (Tuj1) and dendritic (MAP2) markers, (iii) the increase in expression levels of pre-synaptic (Syn1) and post-synaptic (PSD-95) markers upon neuronal maturation, and (iv) the presence in expression of VGLUT1/2 (glutamatergic neurons) and absence in expression of VGAT (GABAergic neurons). Moreover, we verified that the neurons were able to recapitulate basal accumulation of Aβ clusters or phosphorylated tau. Nestin, Tuj1 and MAP2 data are represented as the sum of the neuronal area stained positive for each marker. The expression levels of Syn1, PSD-95, VGLUT1/2, VGAT, Aβ and phospho-tau are reported as the total area or total spot count measured within axonal projections or dendrites, normalised respectively by the total Tuj1 or MAP2 area. **(B)** Representative pictures showcasing the expression levels of each marker at DIV14. Scale bar = 100 µm.

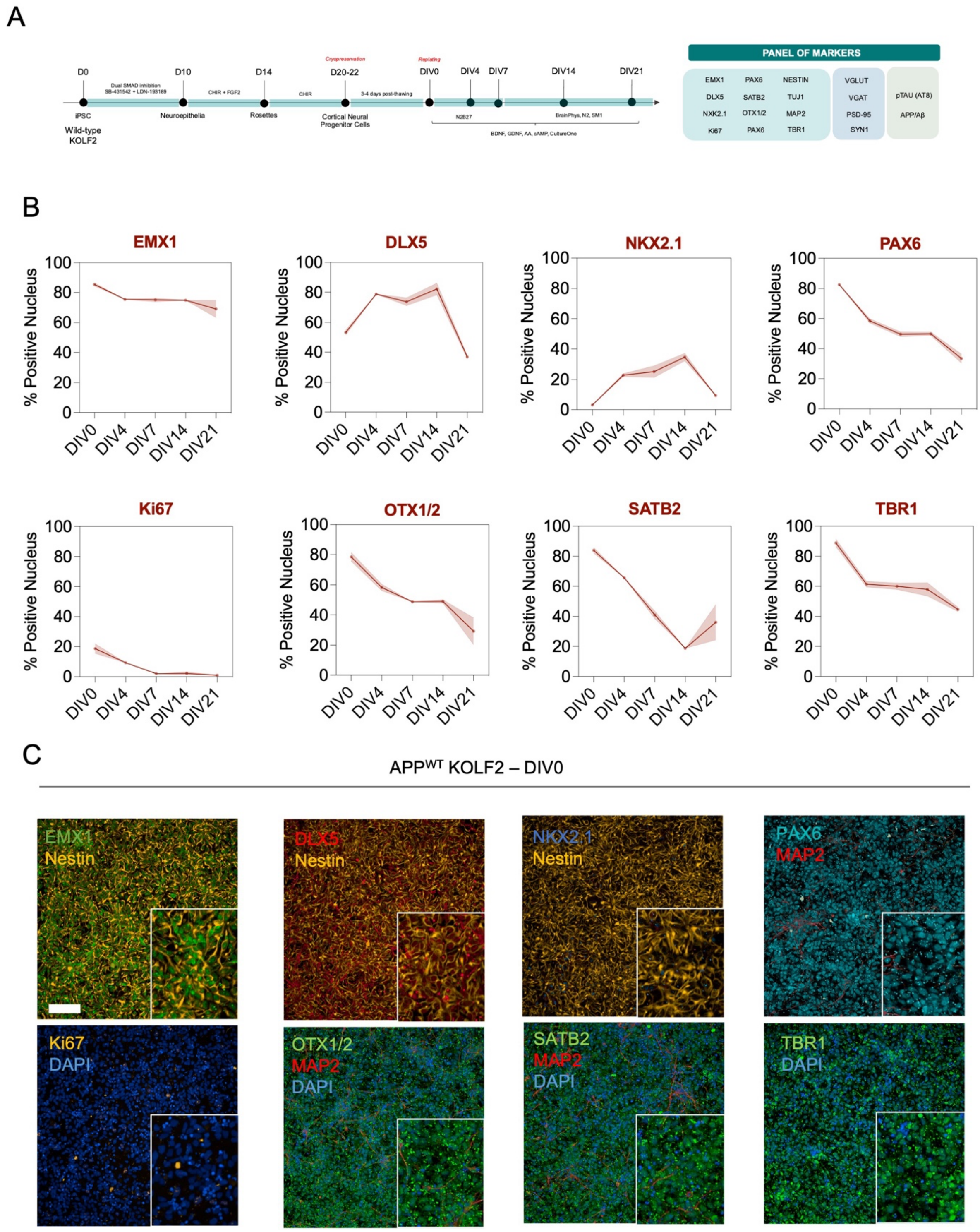


**Figure S3. Quality control of APP$^{WT}$ cortical neural progenitor cells (NPCs) by immunocytochemistry. (A)** Illustration of the differentiation protocol used in this work, which was adapted from a previously reported one (*2*). hiPSCs are converted into neuroepithelia, followed by neural tube-like rosette formation and subsequent isolation of cortical NPCs which then are differentiated to cortical glutamatergic neurons. NPCs generation process takes 20-22 d. Following generation, high-purity cortical NPCs are cryopreserved in large batches to maintain population homogeneity and reduce experimental variability. **(B)** The phenotypic nature of cortical APP$^{WT}$ NPCs is characterised by assessing their enrichment in EMX1, DLX5, PAX6, OTX1/2 and SATB2 expression, which is expected to decrease upon differentiation to mature neurons. Absence of NKX2.1 levels – a marker of ventral forebrain interneurons progenitors (*39*) - was also indicative of the cortical identity of our NPCs. Ki67 was used as a marker to determine the percentage of mitotic cells in culture. Data are represented as the mean and standard error of the percentage of DAPI-positive nuclei that stained positive for the expression of each transcription factor. Expression was assessed by immunocytochemistry at DIV0 (NPC stage), DIV4, 7, 14 and 21 (n = 3). **(C)** Representative images showcasing the expression levels of each transcription factor at DIV0. Cells were also stained for DAPI (nuclei), nestin (cytoskeleton of progenitor cells) and MAP2 (dendritic marker of neuronal maturity). Scale bar = 100 µm.

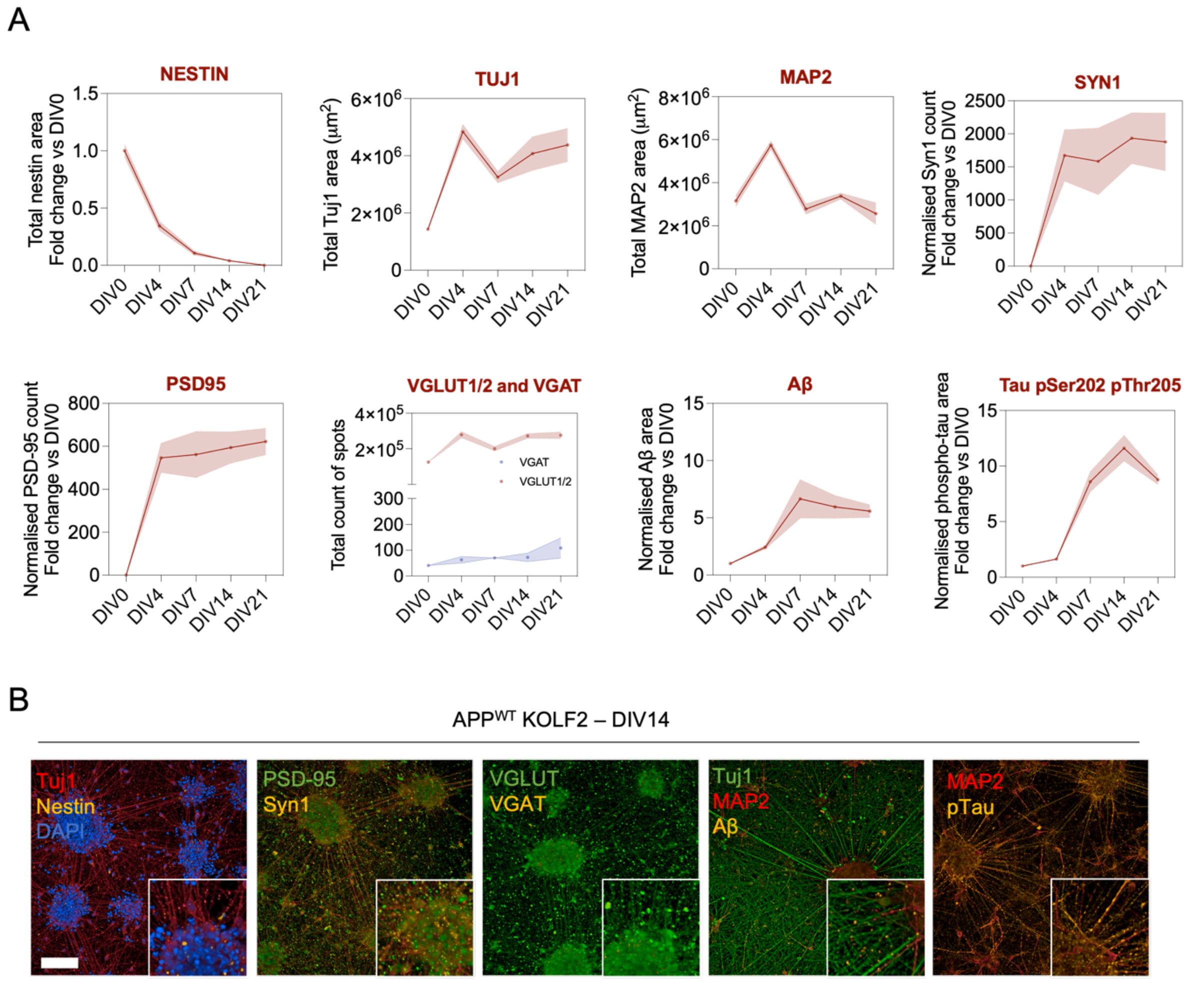


**Figure S4. Quality control of APP$^{WT}$ cortical glutamatergic neurons by immunocytochemistry. (A)** The phenotypic nature of cortical glutamatergic neurons derived from the KOLF2.1 iPSC clone is characterised by assessing: (i) the decrease in expression levels of the progenitor-specific marker nestin upon neuronal maturation, (ii) the expression of axonal (Tuj1) and dendritic (MAP2) markers, (iii) the increase in expression levels of pre-synaptic (Syn1) and post-synaptic (PSD-95) markers upon neuronal maturation, and (iv) the presence in expression of VGLUT1/2 (glutamatergic neurons) and absence in expression of VGAT (GABAergic neurons). Moreover, we verified that the neurons were able to recapitulate basal accumulation of Aβ clusters or phosphorylated tau. Nestin, Tuj1 and MAP2 data are represented as the sum of the neuronal area stained positive for each marker. The expression levelsof Syn1, PSD-95, VGLUT1/2, VGAT, Aβ and phospho-tau are reported as the total area or total spot count measured within axonal projections or dendrites, normalised respectively by the total Tuj1 or MAP2 area. **(B)** Representative pictures showcasing the expression levels of each marker at DIV14. Scale bar = 100 µm.

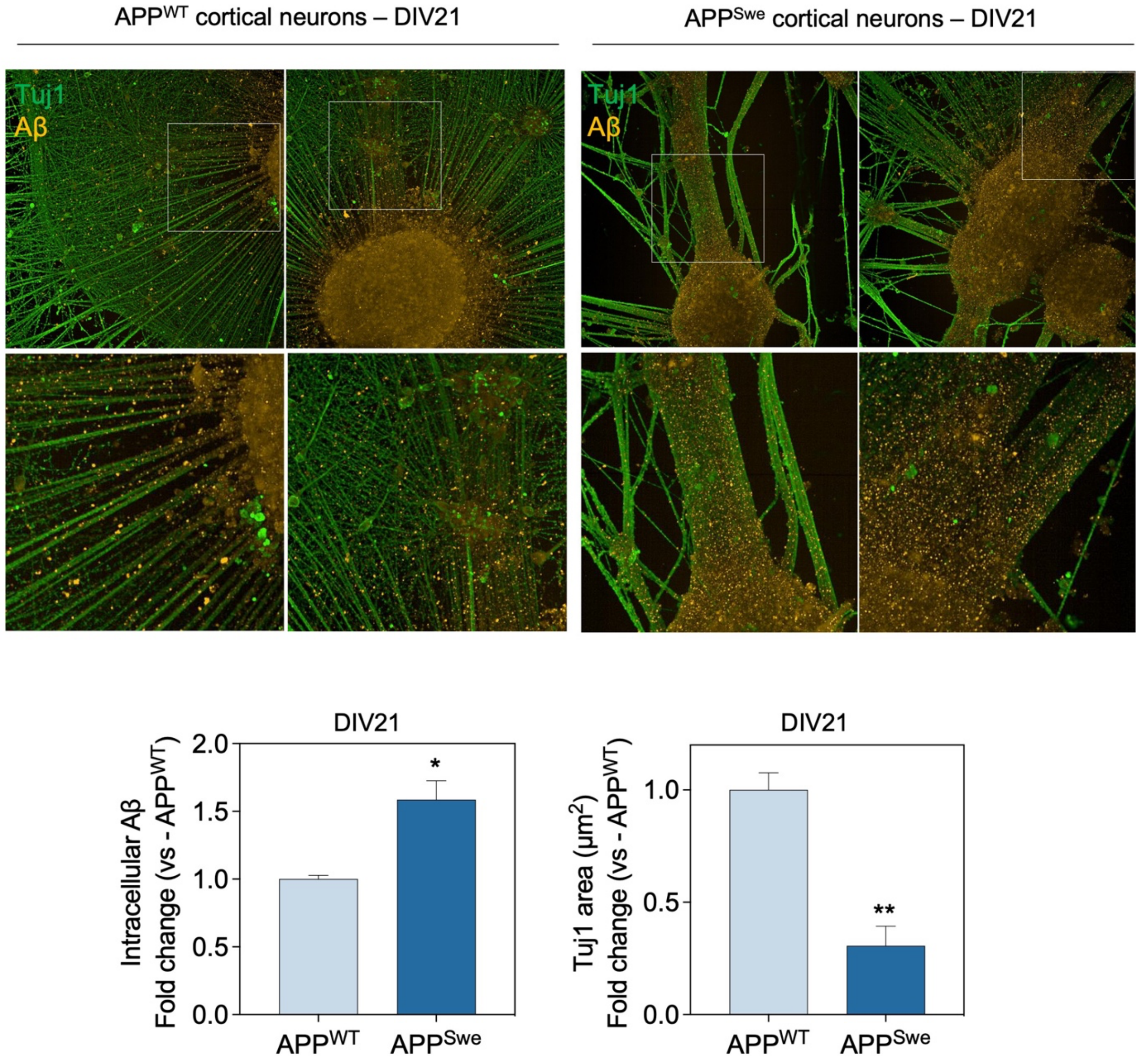


**Figure S5. Differences in axonal distribution and Aβ content between APP$^{WT}$ and APP$^{Swe}$ cortical neurons.** Representative images of APP$^{WT}$ and APP$^{Swe}$ cortical neurons differentiated for 21 d, stained for axonal projections (Tuj1) and Aβ (W02). The images show that APP$^{Swe}$ cortical neurons derived from the clone BIONi010-C-38, tend to form more clusters and thicker axonal projections as compared to the APP$^{WT}$ cortical neurons derived from the KOLF2.1 clone. Therefore, clustering differences cause APP$^{Swe}$ neurons to exhibit a smaller total Tuj1 area not related with degeneration. Levels of total Aβ clusters, measured as the high intensity spots located along the Tuj1-positive area, were 1.5 times higher in APP$^{Swe}$ neurons as compared to APP$^{WT}$. Statistical significance is indicated as: * $p<0.05$, ** $p<0.01$, determined by a t-test.

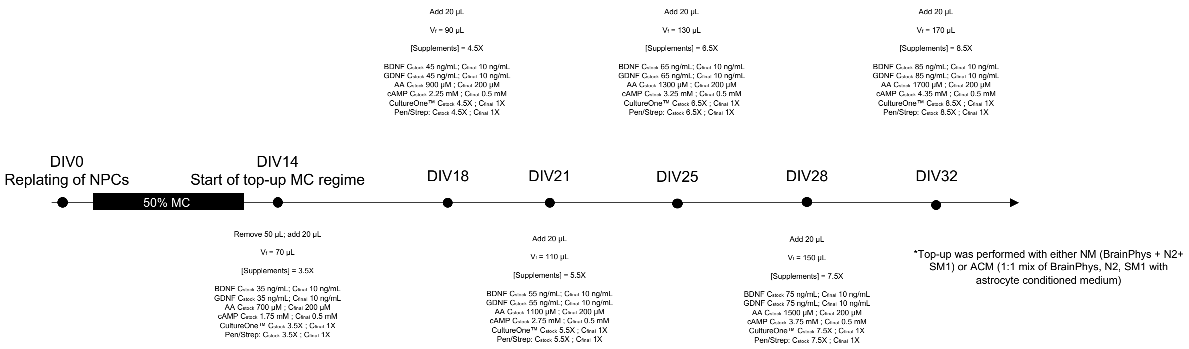


**Figure S6. Experimental pipeline to apply a top-up regime during neuronal maturation.** A 50% medium change regime was applied during the first 14 days of neuronal maturation. At DIV14, the top-up medium change regime was initiated by progressively adding 20 µL per well at the specified days, using progressively concentrated supplemented stocks.

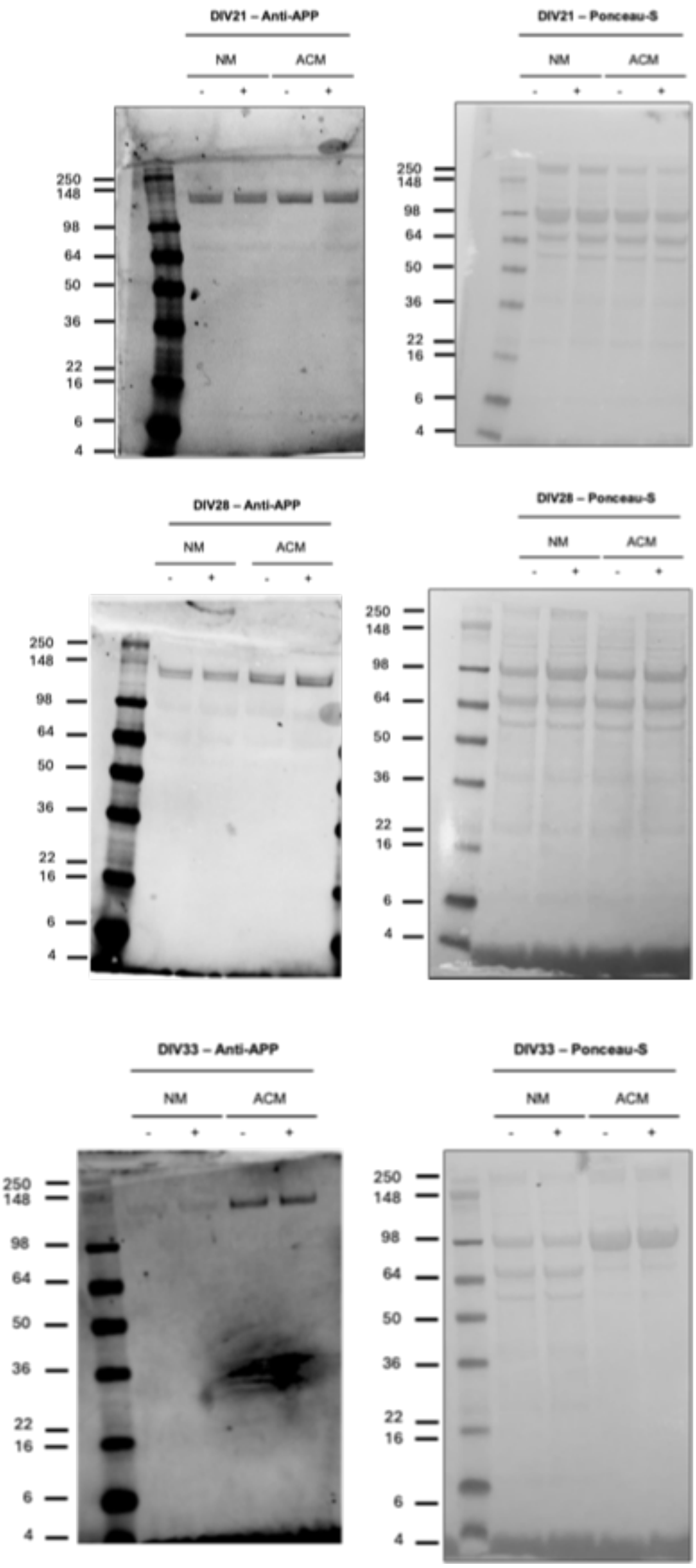


**Figure S7. Western blots for APP and Ponceau-S stainings of NM and ACM treated neurons, pre-treated with ScrRNA (-) or FBXO2 (+) siRNAs**. The antibody W02 was used to measure APP levels.

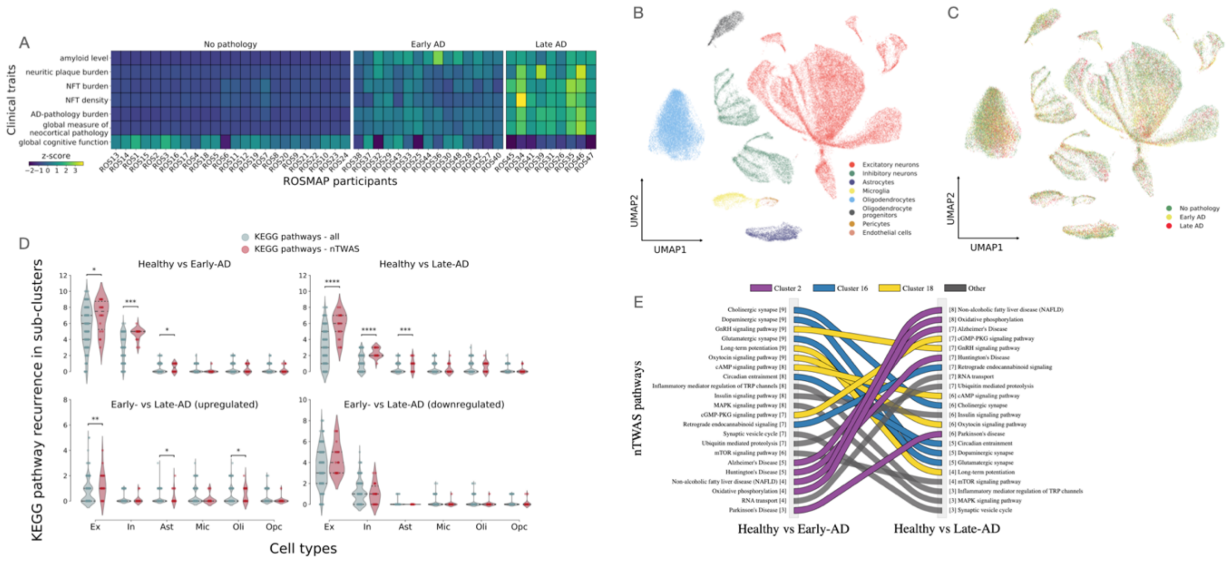


**Figure S7. Bioinformatics analysis of the cell types and biochemical pathways associated with sAD.** (**A**) Grouping of the ROSMAP cohort according to AD clinical readouts. Phenotypic K-means clustering of the 48 individuals from the ROSMAP study (columns), identified by their IDs, using the clinico-pathological observations (rows), measuring the total amyloid level, the neuritic plaque burden, the neurofibrillary tangles burden, and their density, the overall pathological burden, the overall neocortical measure of the pathological state, and the global cognitive function. Z-scored values are shown. (**B,C**) UMAP visualisation of the cell nuclei, with the nuclei coloured according to their cell types. Eight distinct clusters can be identified. (**D**) Among the KEGG pathways, the subset of nTWAS pathways encode the major perturbations occurring in AD in neuronal cells and astroglia. Comparison between the recurrence of the KEGG pathways and its nTWAS subset in all the cell subpopulations in AD stages. The number of times a pathway perturbation was found to be statistically significant within the subpopulations of each cell type (x axes) is reported. The distributions of all KEGG pathways (in grey) and of the nTWAS pathways (in red) are reported for each comparison in each main cell type. The median and interquartile range (IQR) are reported for each distribution with dashed lines. ****$p \leq 0.0001$, ***$p \leq 0.001$, **$p \leq 0.01$, *$p \leq 0.05$, one-sided Mann-Whitney U test with Bonferroni correction for multiple hypothesis testing. **(E)** Sankey plot of the comparison between the recurrence of perturbations in the nTWAS set for the excitatory (Ex) neurons. The nTWAS stage-dependent ranking shows a swap in centrality between signalling and neurodegenerative processes in excitatory neurons. nTWAS pathways, ranked according to the number of subpopulations in which they were found to be dysregulated (number reported next to the pathway names), in the comparison between healthy and early-AD (left side) and late-AD (right-side) individuals, respectively.

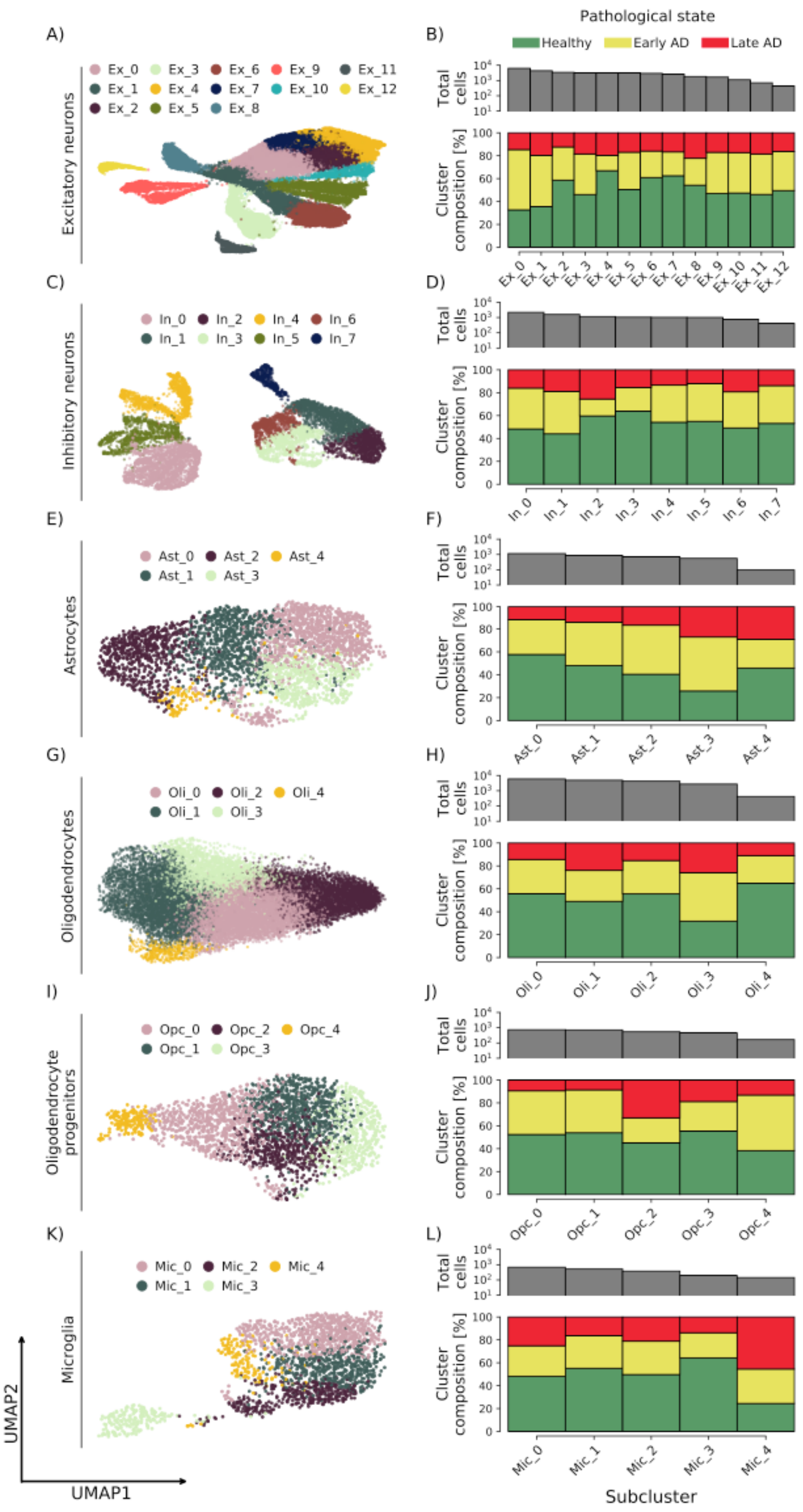


**Figure S8. Louvain graph-clustering generates heterogeneous clusters across the cell types.** Results of the Louvain network-based clustering on the main cell types. **(A,C,E,G,I,K)** UMAP 2D representation of the subpopulations. **(B,D,F,H,J,L)** Clinical composition of the subpopulations, with the total number of cell nuclei belonging to each cluster reported in the top panel.

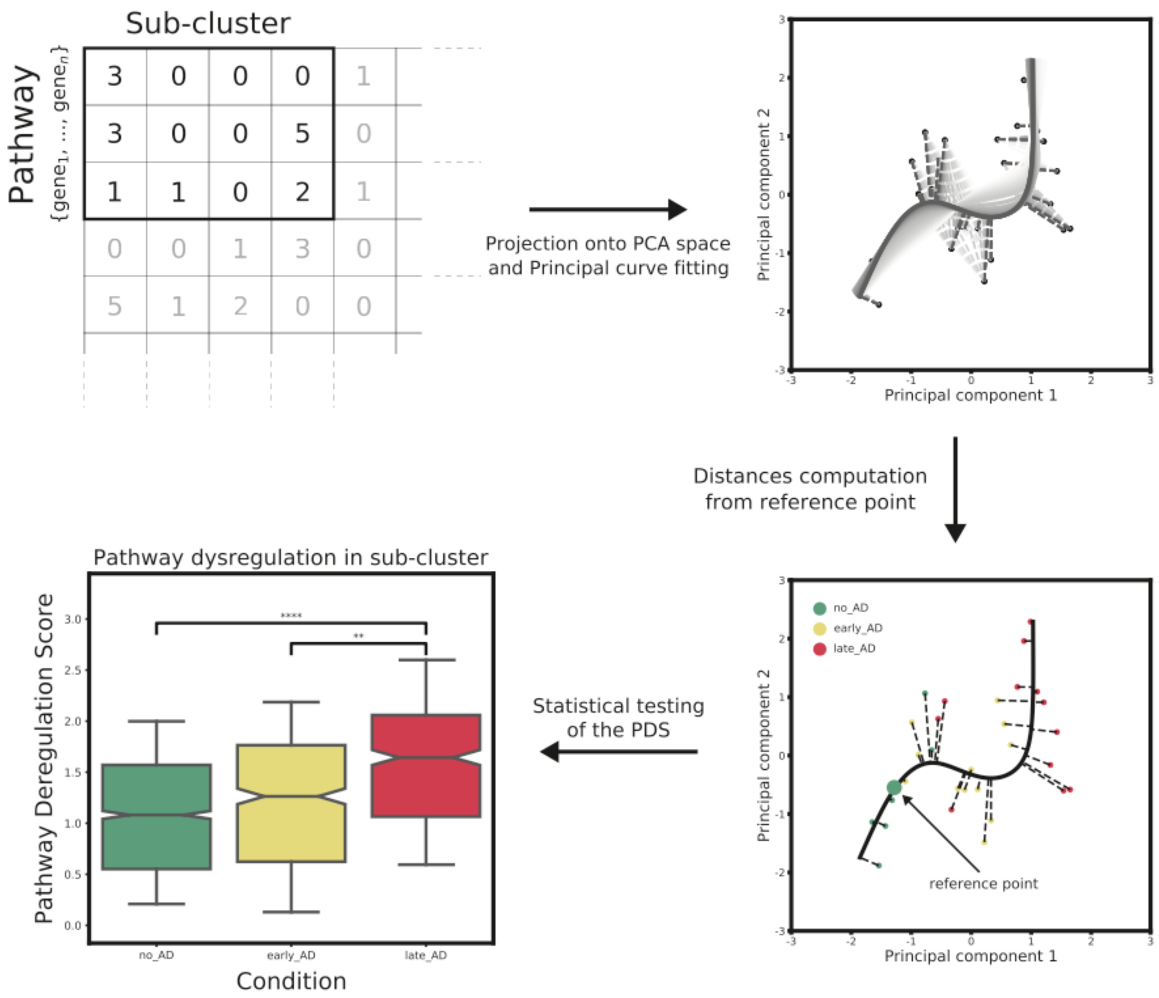


**Figure S9. Pathifier analysis of AD pathways.** To test the perturbation occurring in AD, the gene expression matrix was iteratively transformed into subsets by selecting the matrix corresponding to a specific sub-cluster and a pathway (top left panel). The sub-matrix was then projected onto a lower dimensional space, i.e. the one generated with principal components analysis (PCA), and a principal curve was fitted to the data (the shading shows different iterations of the principal curve calculations to convergence, and the dashed lines the orthogonal projections of the samples onto the iterated curves) (top right panel). The Euclidean distances from a reference point (i.e. the average of the normal samples) of all the samples were then computed (with the colour indicating the pathological state associated with each sample), called pathways deregulation scores (PDS) (bottom right panel). Ultimately, a statistical test was performed to test whether the pathway could separate samples associated with different pathological conditions along the principal curve (bottom left panel).

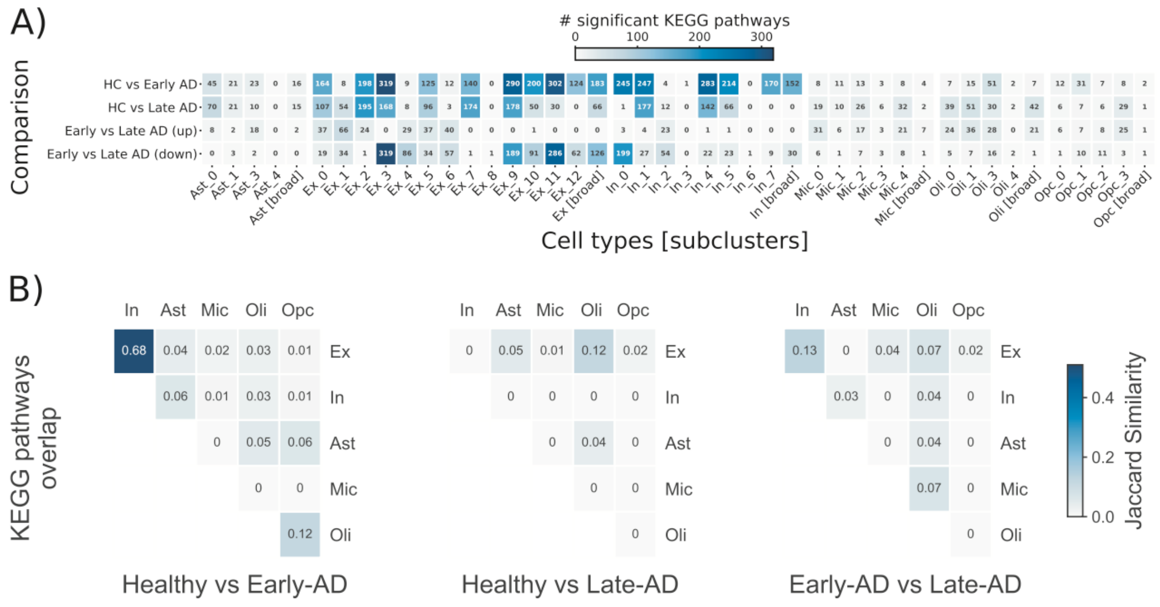

**Figure S10. Cell types show differential and stage-dependent vulnerability to pathways perturbations in AD**. **(A)** Total number of KEGG pathways perturbed in AD in all cellular subpopulations and in the main cell types. Each condition (row) was tested in each subcluster (column). The significance across sub-clusters pertaining to the same cell type was then combined to a determine the number of pathways whose perturbation were considered significant to the cell type as a whole (global cell-type columns are indicated as [broad]). Uprergulation (up) or downregulation (down) tested for the comparison between early and late AD stages is reported as separate. The number of significant pathways is annotated inside each box. **(B)** Overlap of the significant KEGG pathways dysregulated in the cell-types (defined as [broad] in panel A) for each comparison between pathological conditions. Upregulated and downregulated pathways in Early AD (fAD) vs Late AD (sAD) comparison in panel A are merged. The Jaccard similarity is reported inside each box, measuring the fraction of shared pathways.

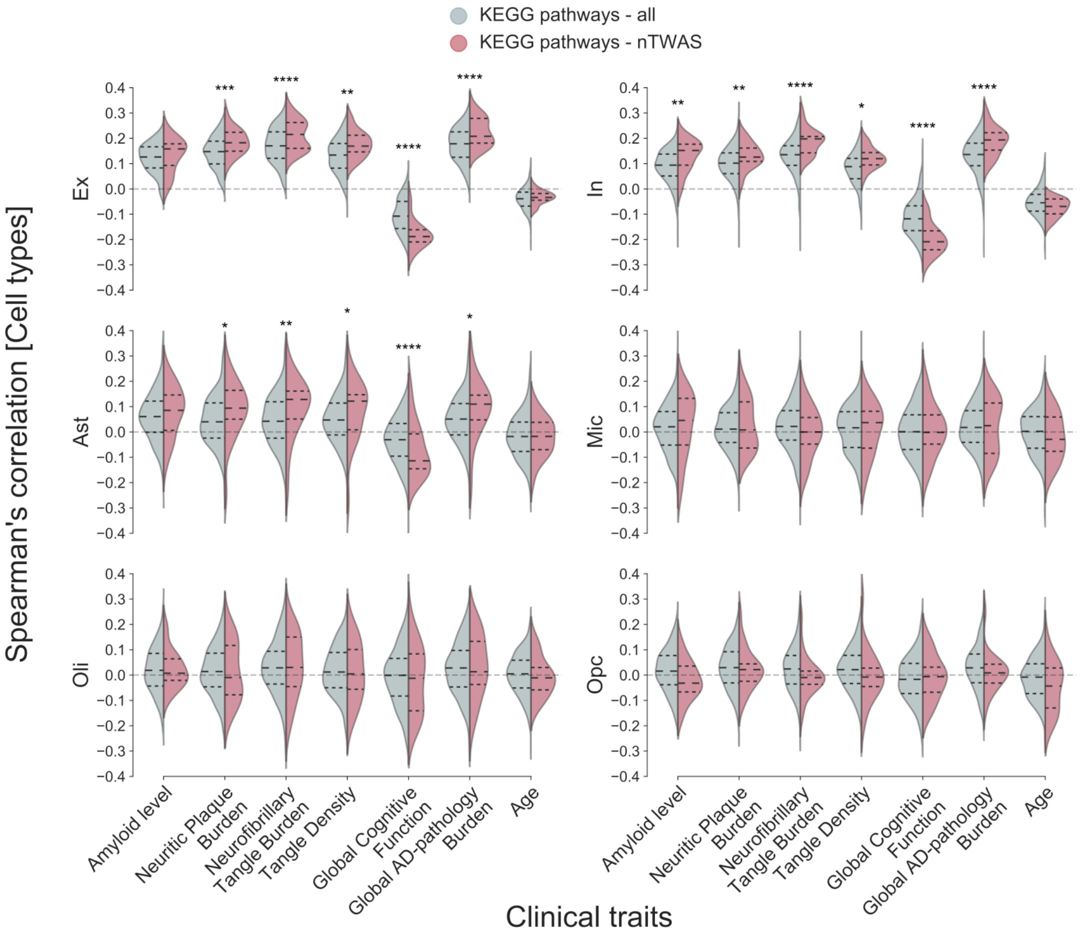


**Figure S11. nTWAS pathways correlate with AD pathological manifestations in neurons and astroglia.** Distributions of the median Spearman's correlation coefficients in the main cell types between the KEGG pathways (grey) and its nTWAS subset (red) and the clinico- pathological traits, reported on the x-axes. The median and interquartile range (IQR) are reported for each distribution with dashed lines. ****p ≤ 0.0001, ***p ≤ 0.001, **p ≤ 0.01, *p ≤ 0.05, one-sided Mann-Whitney U test with Bonferroni correction for multiple hypothesis testing.

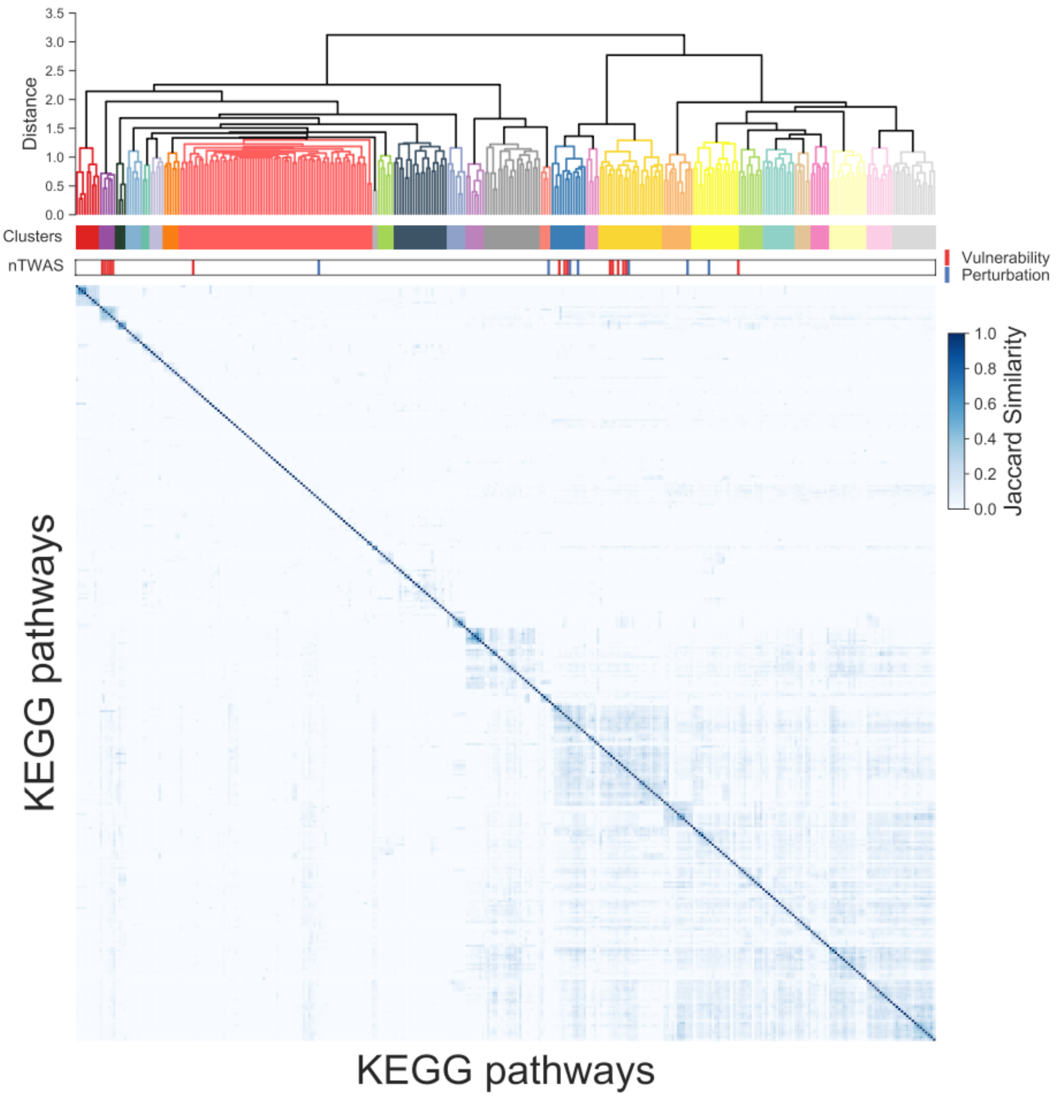


**Figure S12. nTWAS pathways organise in specific clusters within KEGG annotations.** Hierarchical clustering of the KEGG pathways based on the fraction of genes shared amongst them. The dendrogram (top) shows the distances between the clusters. The similarity (bottom) matrix shows the fraction of shared genes measured by the Jaccard similarity. Position of the nTWAS pathways within the clusters (middle) is showed by the vertical bars, with those encoding for the vulnerability of healthy brains to AD identified by the red bars, and those associated with the perturbations occurring in disease reported in blue.

| Cluster | KEGG identifier |
|---|---|
| 1 | hsa00982, hsa00980, hsa05204, hsa00983, hsa00830, hsa00140, hsa00040, hsa00053, hsa00860 |
| 2 | **hsa00190**, **hsa05012**, **hsa05016**, **hsa05010**, **hsa04932**, hsa04714 |
| 3 | hsa05414, hsa05410, hsa05412, hsa04260 |
| 4 | hsa03460, hsa03440, hsa03030, hsa03430, hsa03420, hsa03410 |
| 5 | hsa00350, hsa00360, hsa00400 |
| 6 | hsa00250, hsa00220, hsa00471, hsa04964, hsa00910 |
| 7 | hsa00515, hsa00514, hsa00533, hsa00601, hsa00603, hsa00604 |
| 8 | hsa00230, hsa00240, hsa00760, hsa03010, hsa03008, **hsa03013**, hsa03015, hsa03018, hsa03040, hsa00532, hsa00534, hsa00480, hsa00430, hsa00120, hsa04146, hsa04141, hsa00510, hsa03060, hsa04742, hsa04080, hsa04740, hsa04744, hsa00232, hsa00130, hsa00440, hsa00750, hsa00512, hsa00472, hsa00563, hsa00785, hsa03020, hsa03022, hsa03050, hsa04130, hsa04614, hsa02010, hsa05340, hsa03450, hsa04979, hsa04977, hsa00100, hsa00780, hsa04974, hsa04978, hsa04950, hsa04330, hsa05206, hsa05202, hsa04520, hsa04350, hsa04360, hsa04144, **hsa04120**, hsa04340, hsa04710, hsa04392, hsa00790, hsa00730, hsa00770, hsa00290, hsa00740, hsa00450, hsa00270, hsa00970, hsa04122, hsa00920, hsa03013, hsa04142, hsa00531, hsa00511, hsa00600, hsa01040, hsa00062 |
| 9 | hsa04070, hsa00562 |
| 10 | hsa00051, hsa00520, hsa00030, hsa00500, hsa00052, hsa00524 |
| 11 | hsa04216, hsa00061, hsa00071, hsa03320, hsa00650, hsa00072, hsa00900, hsa00340, hsa00410, hsa00330, hsa00640, hsa00280, hsa00380, hsa00310, hsa00010, hsa00620, hsa00020, hsa00260, hsa00630, hsa00670 |
| 12 | hsa00561, hsa00564, hsa04975, hsa00592, hsa00591, hsa00565, hsa00590 |
| 13 | hsa04672, hsa05310, hsa04940, hsa05330, hsa05332, hsa05320, hsa05416 |
| 14 | hsa04060, hsa04630, hsa04659, hsa04658, hsa05321, hsa05143, hsa05144, hsa05134, hsa05133, hsa05145, hsa05140, hsa05152, hsa04650, hsa04612, hsa05168, hsa04145, hsa05323, hsa04640, hsa05150, hsa04610, hsa05020 |
| 15 | hsa05110, hsa04966, hsa05120, **hsa04721** |
| 16 | hsa05032, hsa04727, **hsa04723**, hsa05033, **hsa04724**, **hsa04713**, **hsa04728**, hsa04726, hsa04926, **hsa04725**, hsa04915, hsa04371, hsa04062 |
| 17 | hsa05030, hsa05031, hsa04962, hsa05034, hsa05322 |
| 18 | hsa04922, hsa04924, hsa04261, **hsa04022**, **hsa04024**, hsa04020, **hsa04921**, hsa04270, hsa04611, **hsa04750**, **hsa04912**, **hsa04720**, hsa04730, hsa04540, hsa04928, hsa04927, hsa04925, hsa04911, hsa04918, hsa04971, hsa04970, hsa04972, hsa04961, hsa04976 |
| 19 | hsa04934, hsa04916, hsa04390, hsa05217, hsa04310, hsa05226, hsa05224, hsa05225, hsa04550, **hsa04150**, hsa05205 |
| 20 | hsa04913, hsa04923, hsa04213, hsa04211, hsa04931, **hsa04910**, hsa04152, hsa04920, hsa04066, hsa05230, hsa04930, hsa04973, hsa04960, hsa04919, hsa04136, hsa04140, hsa04137, hsa04919 |
| 21 | hsa04014, hsa04015, **hsa04010**, hsa04510, hsa04512, hsa04810, hsa05165, hsa04151, hsa05200 |
| 22 | hsa04210, hsa04215, hsa04115, hsa05014, hsa05222, hsa05146, hsa04071, hsa04933, hsa05418, hsa04657, hsa04668, hsa04064 |
| 23 | hsa04218, hsa05166, hsa04110, hsa05203, hsa04114, hsa04914 |
| 24 | hsa05132, hsa05131, hsa05100, hsa05130, hsa04530, hsa04670, hsa04514 |
| 25 | hsa05170, hsa05163, hsa05167, hsa04620, hsa05161, hsa05142, hsa05160, hsa05164, hsa05162, hsa05169, hsa04217, hsa04621, hsa04622, hsa04623 |
| 26 | hsa04072, hsa05231, hsa04664, hsa04370, hsa04666, hsa05235, hsa04660, hsa04662, hsa04625, hsa04380 |
| 27 | hsa05215, hsa05221, hsa04068, hsa04917, hsa04722, hsa04012, hsa05211, hsa05214, hsa05223, hsa05218, hsa05220, hsa05212, hsa05219, hsa05210, hsa05213, hsa05216 |

**Figure S13. List of clusters of KEGG pathways.** The clusters (colour-coded) and the KEGG identifiers corresponding to the pathways contained in each cluster. nTWAS pathways are in bold; hsa00190: Oxidative phosphorylation; hsa05012: Parkinson's Disease (PD); hsa05016: Huntington's Disease (HD); hsa05010: Alzheimer's disease (AD); hsa04932: Non-alcoholic fatty liver disease (NAFLD); hsa03013: RNA transport; hsa04120: Ubiquitin mediated proteolysis; hsa04721: Synaptic vesicle cycle; hsa04723: Retrograde endocannabinoid signaling; hsa04724: Glutamatergic synapse; hsa04713: Circadian en- trainment; hsa04728: Dopaminergic synapse; hsa04725: Cholinergic synapse; hsa04022: cGMP-PKG signaling pathway; hsa04024: cAMP signaling pathway; hsa04921: Oxytocin signaling pathway; hsa04750: Inflammatory mediator regulation of TRP channels; hsa04912: GnRH signaling pathways; hsa04720: Long-term potentiaion; hsa04150: mTOR signaling pathway; hsa04910: Insulin signaling pathway; hsa04010: MAPK signaling pathway. The clusters highlighted in black are enriched in nTWAS pathways.

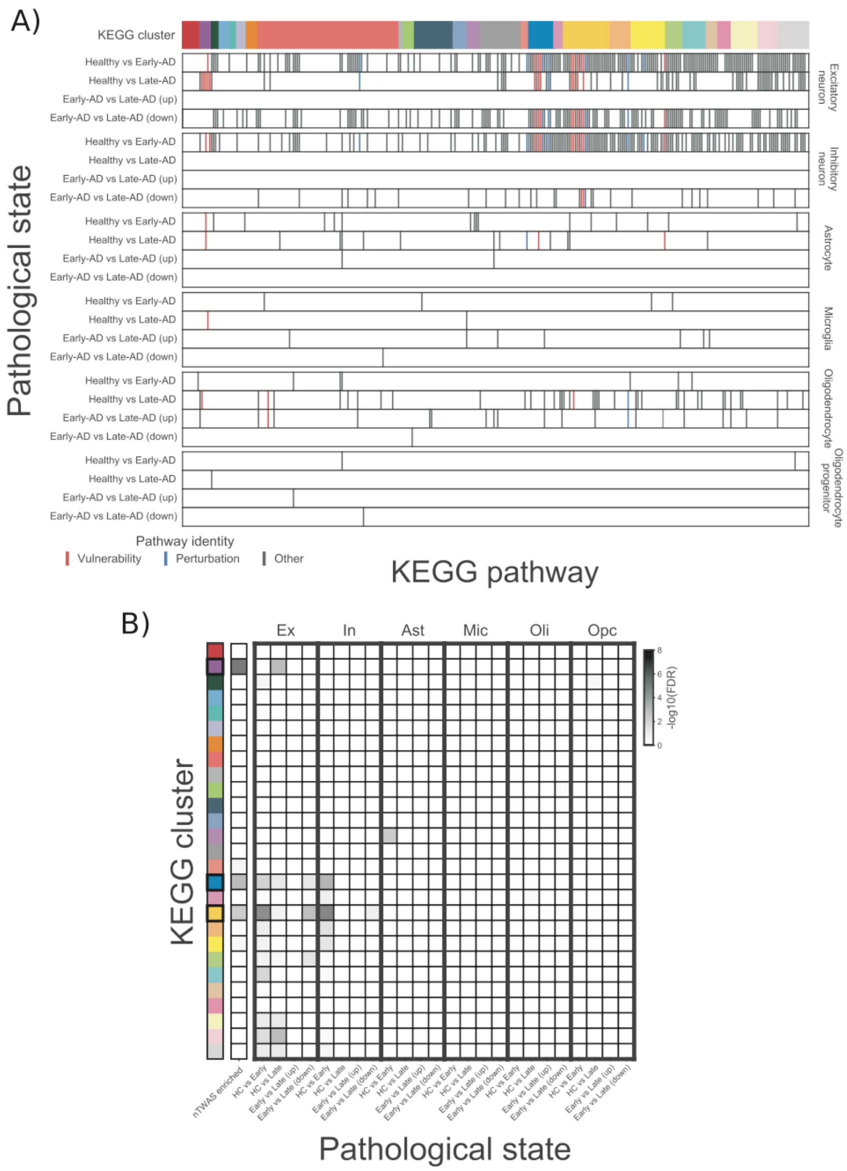


**Figure S14. Clusters of KEGG pathways containing the nTWAS pathways are preferentially perturbed in neuronal cells. (A)** Dysregulated pathways belonging to each KEGG cluster (top), are reported for each cell type (right side of the y axis), and for each comparison within the cell types (y axis). Each vertical bar in the plot corresponds to a pathway, with those belonging to the nTWAS group reported in red (i.e. those encoding for the vulnerability of the brain to AD), and blue (i.e. those found to be perturbed in disease in bulk), while others are reported in black. **(B)** Statistical enrichment of the different KEGG clusters (y axis) in the different conditions (rows), within each cell type (top of the x axis). The first column corresponds to the enrichment of each cluster in nTWAS pathways. Enrichment was computed using Fisher's exact test, with Bonferroni correction for multiple hypothesis testing.
